# Indirect enrichment of desirable, but less fit phenotypes, from a synthetic microbial community using microdroplet confinement

**DOI:** 10.1101/2023.01.11.523444

**Authors:** Ramya Ganiga Prabhakar, Gaoyang Fan, Razan N Alnahhas, Andrew J Hirning, Matthew R Bennett, Yousif Shamoo

## Abstract

Spatial structure within microbial communities can provide nearly limitless opportunities for social interactions and are an important driver for evolution. As metabolites are often molecular signals, metabolite diffusion within microbial communities can affect the composition and dynamics of the community in a manner that can be challenging to deconstruct. We used encapsulation of a synthetic microbial community within microdroplets to investigate the effects of spatial structure and metabolite diffusion on population dynamics and to examine the effects of cheating by one member of the community. The synthetic community was comprised of three strains: a ‘Producer’ that makes the diffusible quorum sensing molecule (*N*-(3-Oxododecanoyl)-L-homoserine lactone, C12-oxo-HSL) or AHL; a ‘Receiver’ that is killed by AHL and a Non-Producer or ‘cheater’ that benefits from the extinction of the Receivers, but without the costs associated with the AHL synthesis. We demonstrate that despite rapid diffusion of AHL between microdroplets, the spatial structure imposed by the microdroplets allow a more efficient but transient enrichment of more rare and slower growing ‘Producer’ subpopulations. Eventually, the Non-Producer population drove the Producers to extinction. By including fluorescence-activated microdroplet sorting and providing sustained competition by the Receiver strain, we demonstrate a strategy for indirect enrichment of a rare and unlabeled Producer. The ability to screen and enrich metabolite Producers from a much larger population under conditions of rapid diffusion provides an important framework for the development of applications in synthetic ecology and biotechnology.

**Abstract Figure:** 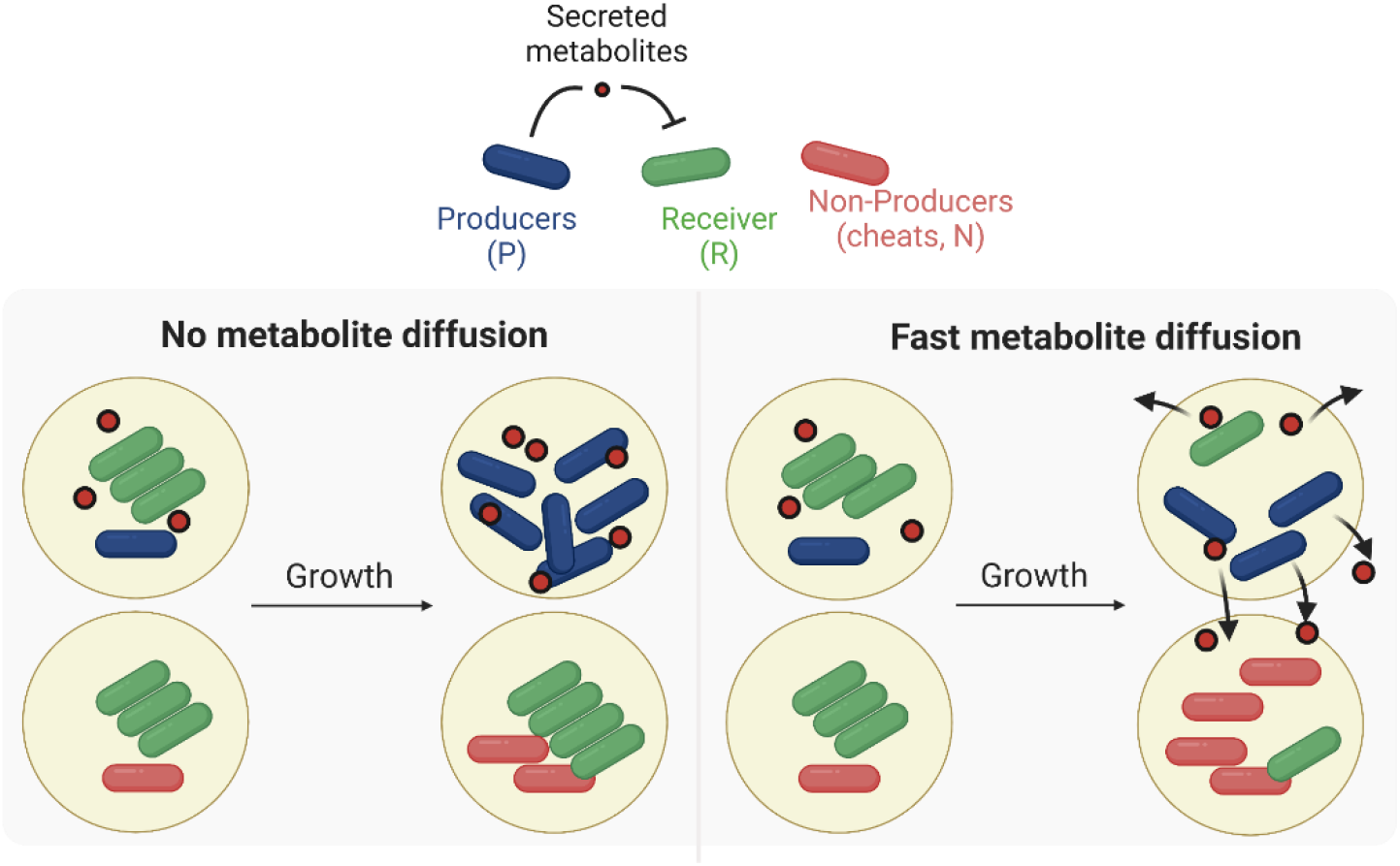

## Introduction

Locey and Lennon estimated microbial biodiversity on Earth to be as high as 1 trillion species^1^. In nature, microbes live in multispecies communities where they compete or cooperate in the utilization of resources within a localized and often structured environment^2,3^. Microbial communities comprise the predominant social structures of life and are critical players in our global ecosystems, yet their potential as contributors to a sustainable future remains largely untapped. Social microbial interactions are dynamic and vary in response to a variety of environmental cues^2,4^. Factors such as spatial patterning, resource availability, and metabolite diffusion are major drivers of microbial community structure across ecological contexts^2,5,6^. Understanding the role of metabolites both as resources, and often as signals, within microbial communities is challenging especially when diffusion plays a significant role in shaping the population dynamics of the community. A quantitative understanding of how diffusion within structured systems affects population dynamics can significantly advance the use of synthetic ecologies for ecological, evolutionary, or biomedical applications^7–10^.

Recent studies of natural microbial interactions have provided key insights into the role of community structure, metabolites, and metabolic networks^11,12^. Such studies can be difficult because of the complexity, inaccessibility, and difficulty of genetic manipulation of highly complex natural communities such as those found in the gut, soil, or aquatic microbiomes^13,14^. Therefore, to provide more tractable experimental approaches for quantitative modeling of community interactions, researchers have begun to use synthetic biology to produce synthetic ecologies using model organisms such as *Escherichia coli* ^5,10,15–18^. Using synthetic biology tools, studies are developed to resemble key features of natural ecosystems such as competition or predator-prey interactions in terms of logic and dynamics^5,19^. Such model systems can be used to develop predictive models and principles to explore the ecological, functional, and key structural features of a community^16,20^.

To assess how diffusible molecular signals, such as AHL, affect population dynamics, we constructed a synthetic community of three competing *E. coli* strains: 1) a ‘Producer’ that synthesizes and secretes AHL; 2) a ‘Non-Producer’ that does not synthesize or respond to AHL, and 3) a ‘Receiver’ that dies in the presence of AHL. Here, the Producer incurs the fitness cost associated with producing the AHL^21^. In a well-mixed environment such as test tubes, the molecules secreted by the Producer are shared by all the members of the population through diffusion, creating an opportunity for cheating by the Non-Producers who do not pay the cost of production, but benefit from the reduced Receiver population^21,22^. However, in a structured environment like microdroplets with reduced permeability of AHL signals, cheating by Non-Producers encapsulated in other nearby microdroplets is minimized. Thus, Producers can preferentially benefit by killing the Receiver despite the cost. Bachmann and colleagues demonstrated a successful use of spatial segregation to identify more efficient, but slower growing variants of *Lactococcus lactis* by serial propagation of the cultures in microdroplets^23^.

To better understand the population dynamics of a community subject to social interactions affected by diffusible metabolites, we encapsulated synthetic communities comprised of the engineered Producers, Non-Producers, and Receivers under various initial conditions. We also set out to identify conditions where a less fit Producer strain that is not labeled could be recovered from a much larger population (which we model as Non-Producers). Indirect Producer enrichment within the context of fast diffusion can be useful for studies in which investigators seek to enrich for wild/unlabeled strains as Producers of valuable biomolecules like antimicrobials.

Our results demonstrate that in the presence of AHL, we observe an increase in the Producer population at the expense of Receivers that die allowing access to more nutrients by the Producers. However, as the number of Producers increases, so does the secreted AHL. The diffusible AHL favors the rise of Non-Producers, which benefit from the metabolites at no cost to their fitness. We developed a mathematical model to examine how different levels of effective diffusion across the microdroplets affects the rate of Producer enrichment. Guided by the modeling results and using precisely controlled bottom-up experiments of simple communities, we also evaluated the effect of initial population ratios, concentrations of metabolite produced, and duration of incubation on population dynamics. Finally, we propose a microfluidic platform incorporating sorting to enrich a rare Producer population secreting diffusible metabolites within a larger mixed community. Insights from this research provide design principles for community studies with unknown or varying degrees of metabolite diffusion in structured environments and guide in screening for rare subpopulations of microbes in both natural and engineered communities.

## Results

### Construction of a spatially segregated synthetic three-strain *E. coli* community that allows diffusion of a molecular signal which impacts the success or failure of all members

To study the dynamics of microbial interactions with diffusible metabolites, we engineered a three-member competitive *E. coli* community consisting of Producer (P), Non-Producer (N), and Receiver (R) strains whose interactions are mediated by AHL (*N*-(3-Oxododecanoyl)-L-homoserine lactone, C12-oxo-HSL) **(Fig. 1a, b)**. The Producer strain synthesizes AHL when *lasI* encoding the acyl-homoserine synthase responsible for AHL synthesis is induced by isopropyl β-d-1-thiogalactopyranoside (IPTG). Once synthesized, AHL can diffuse out of the cell into the media and nearby cells^24^. The Receiver strain contains a plasmid (RG03) that produces constitutive expression (P_Iq_) of the AHL quorum sensing receptor LasR. Together, AHL and LasR activate the promoter (*P_las-lac_*) for *holin* and *lysin* expression leading to cell lysis and death. The Non-Producer strain is a phenotypic and genotypic variant of the Producers that no longer synthesize sufficient AHL to activate the suicide circuit of the Receiver cells. Non-Producers contain the same plasmid (C162b) coding for the IPTG-induced *lasI* gene as the Producers, however, a stop codon has been introduced at the fourth codon of the *lasI* gene (RG01) **(Fig. S1)**. Each strain constitutively expresses different fluorescent proteins for strain identification and quantitation.

**Figure1:**
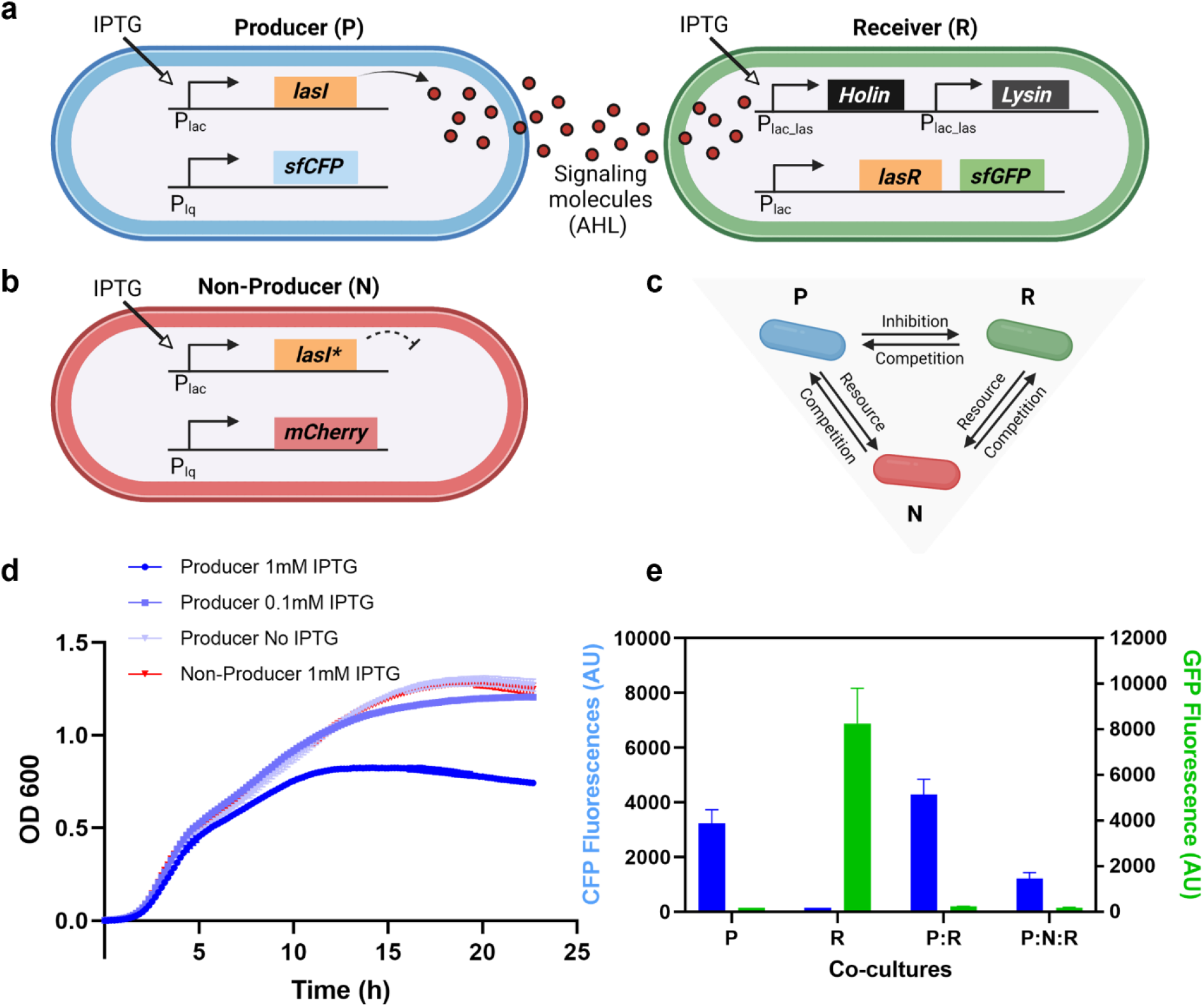
The Producer, the Receiver, and the Non-Producer strain and their social interactions. **a** The Producers secrete AHL to trigger a suicide circuit within Receiver cells, upon induction by IPTG. The secreted AHLs bind to the engineered hybrid promoter (P_las-lac_) along with the constitutively expressed AHL response regulator, LasR to induce the expression of the lysis genes, *holin*, and *lysin* in the Receivers. Expression of these toxic genes within Receivers leads to Receiver cell death. The Producers and the Receivers each produce a different fluorescent protein, *sfCFP* and *sfGFP*, respectively, for strain identification. **b** The Non-Producer strain contains a plasmid coding for IPTG-induced expression (P_*lac*_) of a disrupted *lasI* gene. The Non-Producers constitutively express *mCherry* for strain identification. **c** The Producers can kill the Receiver by producing AHL, but Non-Producers cannot. Producers and Non-Producers themselves compete for resources through exploitative competition. **d** Growth curve for Non-Producers at 1 mM and Producers at different IPTG inductions. (n = 3, error bars = standard deviation (s.d.)) **e** Comparisions of the final population density of the Producers in co-culture with Receivers and Non-Producers, measured using their respective fluorescence proteins. P is the Producer grown in monoculture at 1 mM IPTG after incubation at 37 °C for 16 - 18 hours. R is the population density of Receivers after incubation under identical conditions. P:R is the Producer-Receiver co-cultures, and P:N:R is the Producer-Non-Producer-Receiver cocultures incubated at identical growth conditions with an initially equal population ratio. (n = 3, error bars = s.d.)

We validated the interaction structure of the engineered strains using an agar diffusion assay, where agar pads overlaid with Receiver cells were spotted with AHL or Producers. The Receivers showed very high sensitivity to AHL such that cell death could be observed readily at AHL concentrations as low as 5 nM **(Fig. S2)**. As expected, the Producers can kill the Receivers by the secretion and subsequent diffusion of AHL molecules. All three strains compete for nutrient resources in the shared environment and demonstrate the expected social interaction structure **(Fig. 1c)**.

### AHL biosynthesis imposes a strong fitness cost on the Producer strain

The fitness cost on Producers due to AHL production was measured by plotting the Producer growth curves with different IPTG concentration **(Fig. 1d)**. As shown in **Fig. 1d**, as IPTG concentration increased, the final density of the Producer decreases with increased *lasI* expression. In the absence of IPTG, the calculated growth rate **(Table S1)** and the observed final densities of the Producers are comparable to that of the Non-Producers. When the Producers are induced with 1 mM IPTG, which corresponds to the strongest induction of QS molecule production, Producers have a lower growth rate (0.5404 /h) and reaches a lower final optical density of ~0.8 compared to the Non-Producers (growth rate of 0.6416 /h and final optical density of ~1.3), under the same growth conditions **(Fig. 1d, S3)**. This data suggests there is a fitness trade-off between the costly AHL molecule production and the growth of the Producers.

### Producers rise in frequency in well-mixed Producer/Receiver co-cultures despite the fitness cost, but decline when confronted with a cheating Non-Producer as a competitor

To provide quantitation of each strain during various combinations of co-culturing, each strain was labeled with distinct fluorescent proteins that were expressed constitutively. Fluorescence intensities were recorded and used as a proxy for the density of each strain (Producers: super-folder Cyan Fluorescent Protein (*sfCFP*), Non-Producers: mCherry, and Receivers: super-folder Green Fluorescent Protein (*sfGFP)*). When equal population ratios of Producer-Receiver (P:R) are co-cultured at 1 mM IPTG, the Producers are able to rapidly eliminate AHL-sensitive Receiver competitors and dominate the population **(Fig. 1e, S3)**. In fact, Producers can eliminate 100-fold excess of Receivers at 1 mM IPTG induction (**Fig. S4**). As expected, when the Non-Producers and the Receivers (N:R) are co-cultured in the absence of Producers and under identical conditions, with IPTG, N:R compete comparably for the shared resources **(Fig. S5)**.

When the three strains (P:N:R) are co-cultured in initially equal population ratios at 1mM IPTG, the population density of the Producers is significantly reduced **(Fig. 1e)**, even as the Receiver population declined to extinction. Since Non-Producers do not have a fitness cost associated with AHL production, they can grow faster than Producers in this mixed culture **(Fig. S5)**. Non-Producers therefore cheat successfully by benefiting from the production of AHL by the Producer. Thus, we established a community where the benefit of eliminating competition for the common resources without the fitness cost leads to the rise of the Non-Producing cheaters.

### Mathematical population model of the three strains

To gain quantitative insight into the populational dynamics when co-culturing the three strains, we turn to mathematical modeling. Let *P*(*t*), *N*(*t*), and *R*(*t*) represent the population count of the Producers, Non-Producers, and Receivers, correspondingly at time *t*. In a well-mixed environment, we assume AHL signals are uniformly distributed and its concentration is denoted by *Q*(*t*). Assuming logistic growth with a carrying capacity of *S_max_*, we have the following ODE model (see S.I. for more details),

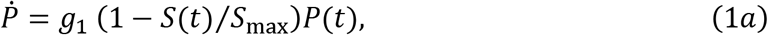

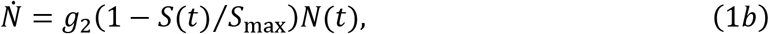

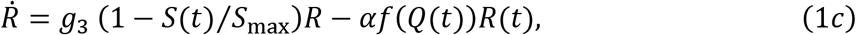

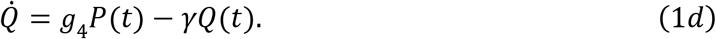

Here, we set *S*(*t*) = *wP*(*t*) + *N*(*t*) + *R*(*t*), with *w* ≥ 1 and increases with [IPTG], to account for the fitness cost of Producers synthesizing AHL molecules **(Fig. S3)**.

### AHL diffuses quickly among the microdroplets

A well-known strategy to prevent Non-Producers from cheating is by spatially segregating the Producers from the Non-Producers^21,22^. While the privatization of cells within the emulsion microdroplets is well established in the literature, the effect of metabolite diffusion is not as widely studied^25–30^. To examine the privatization of AHL molecules within microdroplets, we co-encapsulated Producers (*λ*_1_ = 0.1), and Receivers (*λ*_2_ = 5), in 40 μm droplets at 1 mM IPTG **(Fig. 2a, S6)**. Here, *λ_k_* represents the average number of cells per microdroplet for the corresponding cell type, for *k* = 1,2,… Because we use flow-focusing microfluidic devices to produce co-encapsulated populations of cells, the distribution of cell counts for each strain within the emulsion of microdroplets is assumed to follow a Poisson distribution^31,32^. The probability of a microdroplet containing *n*_1_ Producers, and *n*_2_ Non-Producers, and *n*_3_ Receivers, can be calculated as follows^33^,

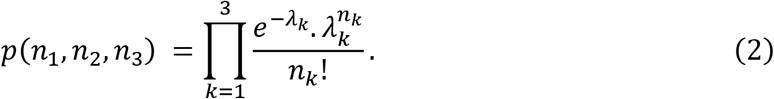

We expect that ~90% of the microdroplets will contain only Receivers, see equation (S3). Since Receivers are tagged with *sfGFP*, if AHL does not diffuse between the microdroplets, we expect that a strong *sfGFP* signal will be present in that 90% microdroplet subpopulation. However, experimentally, we see that no microdroplets contain Receivers within 24 h of growth, suggesting that the AHL molecules secreted by the Producers can diffuse to the neighboring microdroplets and kill all Receivers **(Fig. 2b)**. In fact, when we mixed microdroplets containing AHL molecules, with microdroplets containing AHL reporters, we observed reporter expression within 2 hours of incubation **(Fig. S7, S8)**. AHL molecules are known to partially dissolve in the oil phase as they are slightly hydrophobic and can therefore diffuse directly through the oil and also at the interface between the adjoining microdroplets^25^. However, the diffusion rate in microdroplets is expected to be slower than in well-mixed suspension with a diffusivity of D_oil_ ≈ 1 μm^2^/s in microdroplets and D_bulk_ ≈ 100−1000 μm^2^/s in bulk^25^.

**Figure 2:**
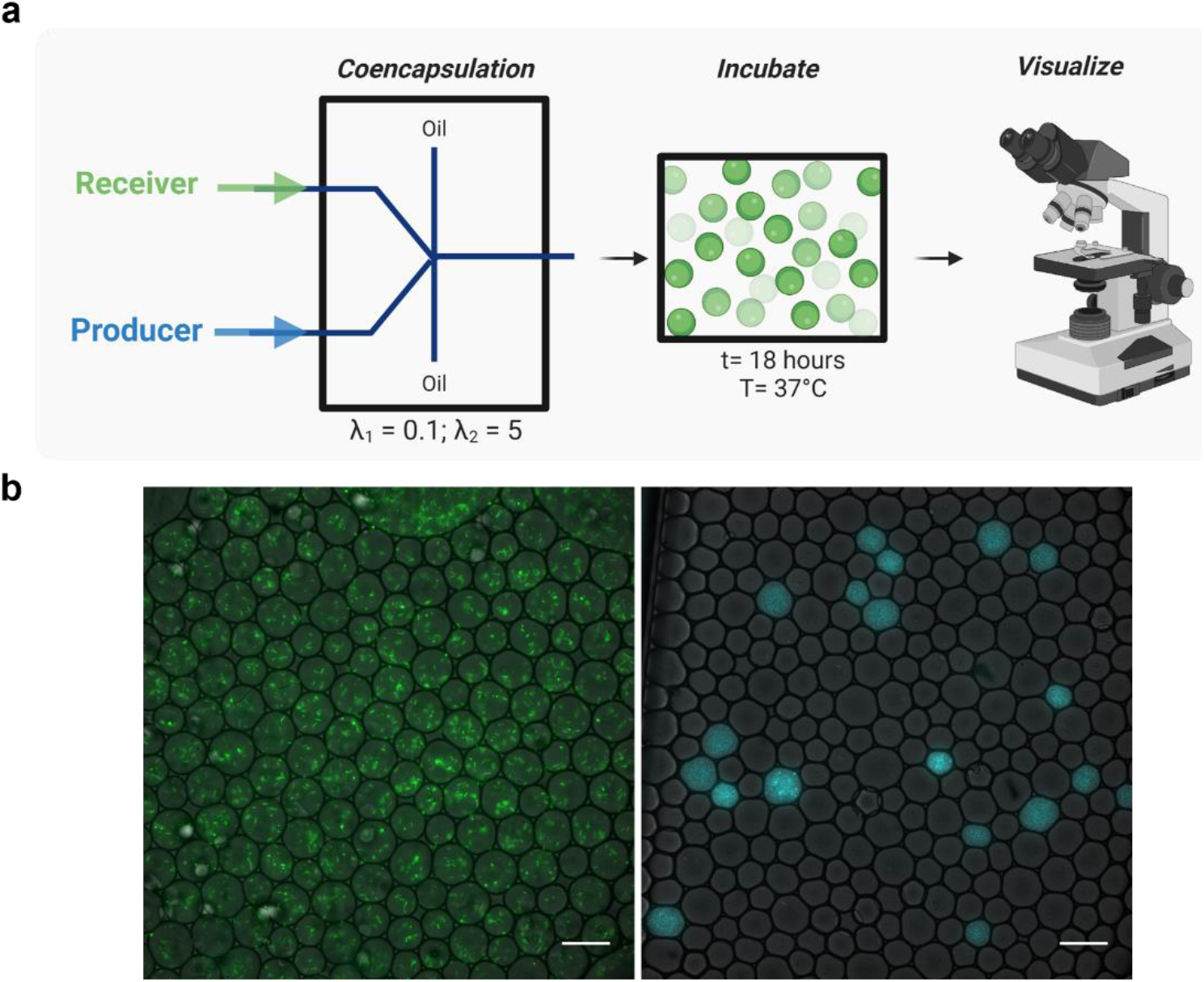
AHL diffuses rapidly between the emulsion microdroplets. **a** Schematic of the experimental setup to test the privatization of AHL molecules within the microdroplets. Producers and Receivers induced by 1 mM IPTG were co-encapsulated within the microdroplets, using the microfluidic droplet generator. The collected microdroplets were incubated for 16 - 18 h at 37 °C. After incubation, the microdroplets are imaged under the microscope. (Scale bar 50 μm. n = 3.) **b** At t = 0 h, due to the higher density of Receivers, all the microdroplets have at least 1 Receiver (*λ*_2_ = 5; green). While less than 10% of the microdroplets have a single Producer (*λ*_1_ = 0.1; white). Since the exposure time for Producers is optimized for the signal at maximum growth, we cannot see single encapsulated cells at t = 0 h. After incubation, as AHL molecules diffuse between the microdroplets, only the Producer cells are observed but all the Receiver cells are killed, even in the neighboring microdroplets.

When modeling population dynamics in a spatially segregated environment, we first characterize microdroplets by the initial population composition of each strain after co-encapsulation. The total population is then calculated as the weighted sum of each type of droplets, indexed by *j*, with the corresponding weight described by equation (2). Let *P_j_*(*t*),*N_j_*(*t*), and *R_j_*(*t*) represent the Producer, Non-Producer, and Receivers population in microdroplet type *j* at time *t*, whose interactions follow Equation (S6a-c). Let *Q_j_*(*t*) denote the AHL levels in the *j^th^* type of microdroplet. Assume that some portion *β* of the signals produced are local to the microdroplet of production while the other 1 − *β* portion is available to all microdroplets. We then have,

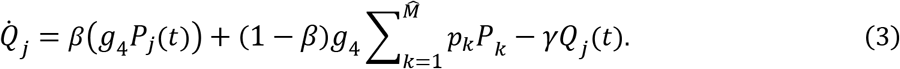

Here, setting *β* = 1 represents the extreme case where AHLs doesn’t not diffuse out of the microdroplets, while setting *β* = 0 corresponds to the other extreme case of fast signal diffusion, where signals produced are immediately shared with all microdroplets. The effective diffusion is impacted by a combination of factors, such as high percentage of microdroplets containing Producers, high Producer population inside the microdroplets, fast synthesis rates and fast diffusivity of AHLs. Modeling the in-between case of *β* ∈ (0,1) is motivated for scenarios when AHL levels in the system is relatively low, where the sequestration of AHL molecules by Receivers plays a considerable role.

In our scenario, Producers with *λ*_1_ = 0.1 leads to ~10% of the microdroplets containing at least one Producer. Assuming a 3D arrangement of the microdroplets, 10% means, on average a microdroplet containing no Producer is right next to a microdroplet with a Producer. Moreover, the timescale of cell growth (hours) is much slower than the timescale of AHL diffusion across to the neighboring microdroplet (seconds). Therefore, we assume that AHL concentrations are the same for all microdroplets in the system.

When the beneficial resources are completely privatized within the microdroplet (non-diffusible between microdroplets), Millet and colleagues showed that muconate enzyme producers are serially enriched 6-fold in microdroplets compared to the well-mixed bulk cultures^34^. Whereas, we have a culturing condition where the cells are spatially segregated, but the secreted AHLs diffuse among microdroplets. Consequently, the Non-Producers, when co-encapsulated with Receivers, can cheat by benefiting from reduced competition, due to AHL diffusion from neighboring Producer microdroplets and killing Receivers. Understanding the effects of diffusible public goods on the population dynamics of the three strains can provide relevant insights into enriching for novel producers in natural systems as well as synthetic microbial communities. Since the AHL molecules diffuse rapidly, we are essentially stress testing the limits of the microdroplet system as culturing platforms to enrich for novel metabolite producers.

### A spatially segregated microdroplet environment improves the Producer population growth in a co-culture even with rapid diffusion, compared to well-mixed suspension

To enrich for Producer cells in co-culture with faster-growing Non-Producers, we designed and tested a competition-based experimental strategy **(Fig. 3a)**. We reintroduced Receivers at each iteration to compete with Non-Producers. The co-cultivation of the three strains was achieved by co-encapsulating the three cultures at chosen densities induced at 1 mM IPTG, in particular, 1.5*10^6^ cells/ml of Producers, 1.5*10^6^ cells/ml of Non-Producers, and 1.5*10^8^ cells/ml of Receivers. This corresponds to starting λ values of Producers *λ*_1_ = 0.05, and Non-Producers *λ*_2_ = 0.05, Receivers *λ*_3_ = 5, respectively (a 1:1:100 starting ratio of P to N to R). After 24 hours of incubation, the cultures were propagated for the next round of growth, diluting 50-fold, and spiking the population with 1.5*10^8^ cells/ml of Receiver (*λ*_3_ = 5), to replenish the Receivers killed in the previous iteration. The populations were iterated every 24 hours for 3 cycles. In parallel, the strains were grown under identical population densities in test tubes to examine a well-mixed batch environment **(Fig. 3b)**.

**Figure 3:**
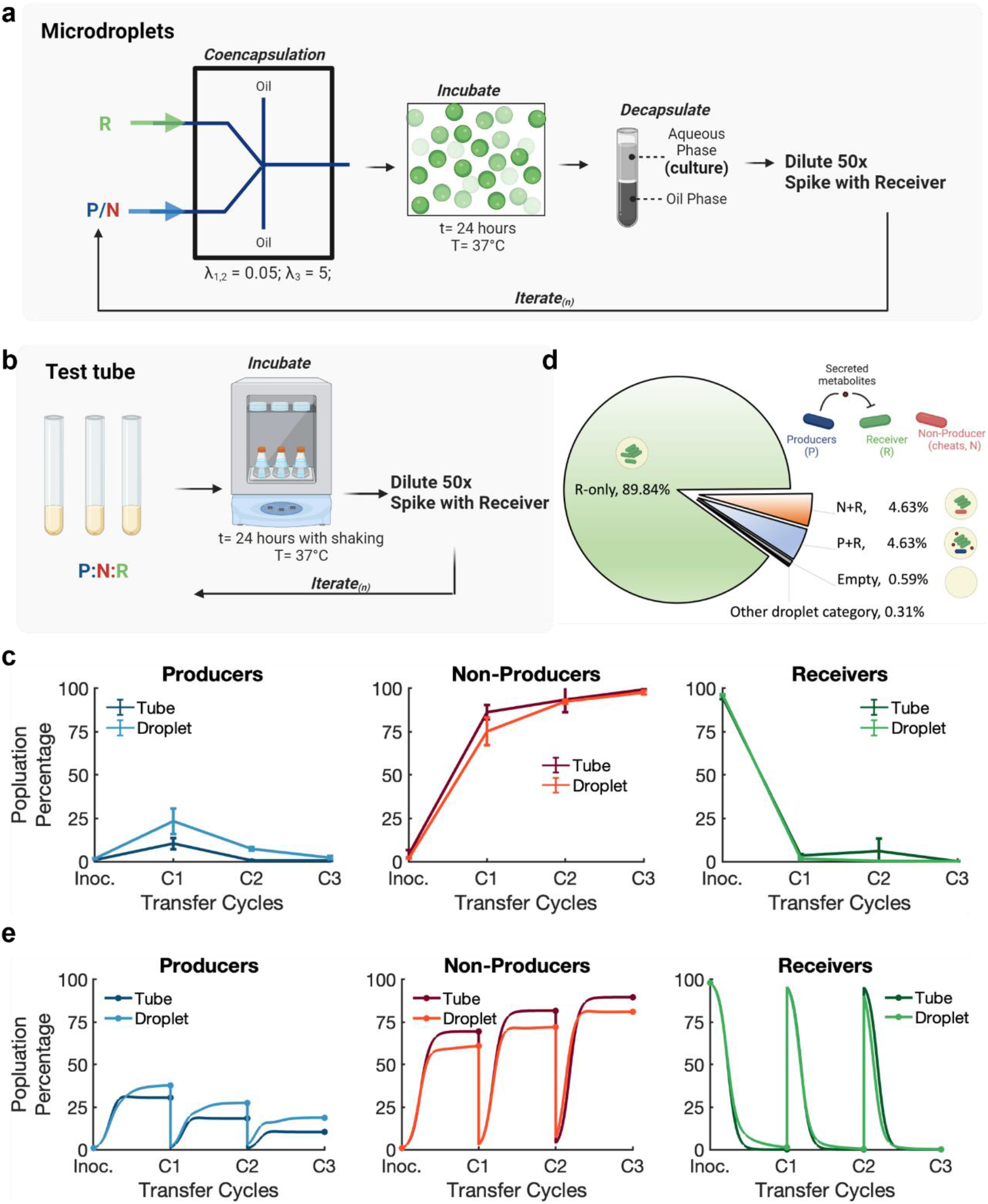
The Producer, the Non-Producer, and the Receiver growth trends in the competition-based experiments in microdroplets versus well-mixed test tube conditions. **a** Schematic of the experimental strategy in microdroplets and well-mixed test tube **b** conditions. Microdroplets were incubated at 37 °C without shaking to avoid droplet coalescence, while the test tubes were incubated at 37 °C with shaking to produce well-mixed conditions. After 24 h, the cultures were diluted 50-fold and spiked with Receivers for the next iteration. **c** The relative cell counts corresponding to Producer, Non-Producer, and Receiver population percentages were used to compare the growth trends of the two culturing conditions. At the end of the first growth cycle, in both the microdroplets and the well-mixed conditions, Receivers went extinct while Producers and Non-Producers grew successful. The Producers grew to a higher percentage (~23%) in the spatially segregated microdroplets system compared to the well-mixed conditions (~10%) (n = 2, error bars = s.d.). **d** Pie chart for the percentages of different categories of microdroplets characterized by the presence of different encapsulated strains. Other droplet *categories* include, P+N+R (~0.24%), P-only (~0.03%), N-only (~0.03%), and P+N (~0.002%). **e** Simulation results of the population percentages for Producer, Non-Producer, and Receiver for the well-mixed test tube conditions described by equations (1) and the mircrodroplet condition described by equations (S6). Parameters used are listed in Table S2.

At the end of the first growth cycle, we observed that, in both the microdroplets and the test tube conditions, Receivers became extinct while Producers and Non-Producers grew successfully. Importantly, a ~2-fold (from about 10% to 23%) higher percentage of Producers were present at the end of the first growth cycle in the spatially segregated microdroplets system than in well-mixed test tube conditions (**Fig. 3c, S9**). We expect that the higher Producer percentage is the result of the spatial segregation of Producers from Non-Producers. In particular, the faster-growing Non-Producers will stop growing once it reaches the carrying capacity of the microdroplets they are in. In the well-mixed environment of the test tubes, Non-Producers can make use of the entire volume of the culture. The effect of spatial segregation on Producers and Non-Producers was also shown by our mathematical modeling (see S.I. for the full microdroplet model description). Upon the initial stochastic encapsulation, Producers and Non-Producers only co-exist in less than 0.24% of the microdroplets. Most of the Producers are co-encapsulated with Receivers only, as shown in **Fig. 3d**. Similarly, due to symmetry with *λ*_1_ = *λ*_2_, most of the Non-Producers are co-encapsulated with Receivers as well. Thus, the spatial segregation of P and N is achieved by having *λ*_1_ ≪ 1 and *λ*_2_ ≪ 1.

During incubation, in the P+R microdroplets, Producers can kill all Receivers and grow to the carrying capacity of the microdroplet. Whereas in the N+R microdroplets, Non-Producers compete with Receivers. The effectiveness of the N:R competition is therefore determined by the effective diffusion of AHL signals. Our calculation suggestion that, with *λ*_2_ = 0.05 for the Producers, we expect approximately 4.9% of the droplets contains the Producers. Moreover, AHL signals can be sequestered in Receiver-only microdroplets, which comprise about 90% of all microdroplets. However, as suggested by **Fig. 3c**, Receivers cells are all killed. Therefore, we use the fast-diffusivity limit described by equation (3) with *β* = 0 to model the distribution of the AHL signals. Under this extreme, where signals are evenly distributed among all droplets, Receivers in the N-R droplets are being killed as fast as the ones in the P+R droplets. This inefficient competition also leads to decreasing Producer population ratio in later sequential incubation cycles, as shown in **Fig. 3c**. The simulation results in the same trend, as shown in **Fig. 3e**. As suggested by the modeling result, to improve the effectiveness of the competition within the microdroplet system, we need to lower the AHL production by reducing the IPTG induction levels and lowering the initial Producer population. In addition, as shown by the experimental results **(Fig. S10)** and confirmed by the simulation using estimated parameters **(Fig. 3e)**, Receivers are mostly killed by the end of 12 hours. Based on these observations, we reduced our cycle length from 24 hours to 12 hours to improve and further validate the efficacy of our Receiver competition strategy for enrichment of Producers.

### Producer population density increases in microdroplets when lower concentrations of AHLs are produced

AHL production is regulated by the exogenously added inducer IPTG **(Fig. S11)**. By controlling the amount of AHL produced through IPTG induction, we can control the extent to which Receivers are killed over time. Here we co-cultured the three strains in emulsion microdroplets induced by two different levels of IPTG concentration: 1 mM, and 0.05 mM. The initial population density was chosen as before with *λ*_1_ = 0.05 for Producers, *λ*_2_ = 0.05 for Non-Producer, and *λ*_3_ = 5 for Receivers, (a 1:1:100 starting ratio of P to N to R). After 12 h of growth, the cell counts for each strain were measured for 6 cycles **(Fig. 4a, S12)**. At the end of the first cycle, we observe that the Producer percentage increased ~7-fold (from about 6.3% to 42%) under the 0.05 mM IPTG inducer condition than in the higher 1 mM IPTG condition. The lower IPTG level leads to a lesser metabolic burden on Producer growth. Moreover, with less AHL produced, there are more Receivers to compete against Non-Producers in the N+P microdroplets. Taken together, decreasing the AHL biosynthesis burden on the Producer coupled with increased competition on Non-Producers led to a higher Producer percentage at the lower IPTG level, as shown by our simulation results **(Fig. 4b)**. As we iterate the growth cycle, we observe that the Producer percentage drops in both conditions **(Fig. 4a)** although much more gradually at the lower IPTG concentration. Interestingly, we observe that, by the end of the 6^th^ cycle, the Receiver percentage starts to rise while the Non-Producer percentage falls, indicative of the increased competition from the Receivers, with Producers mostly gone.

**Figure 4:**
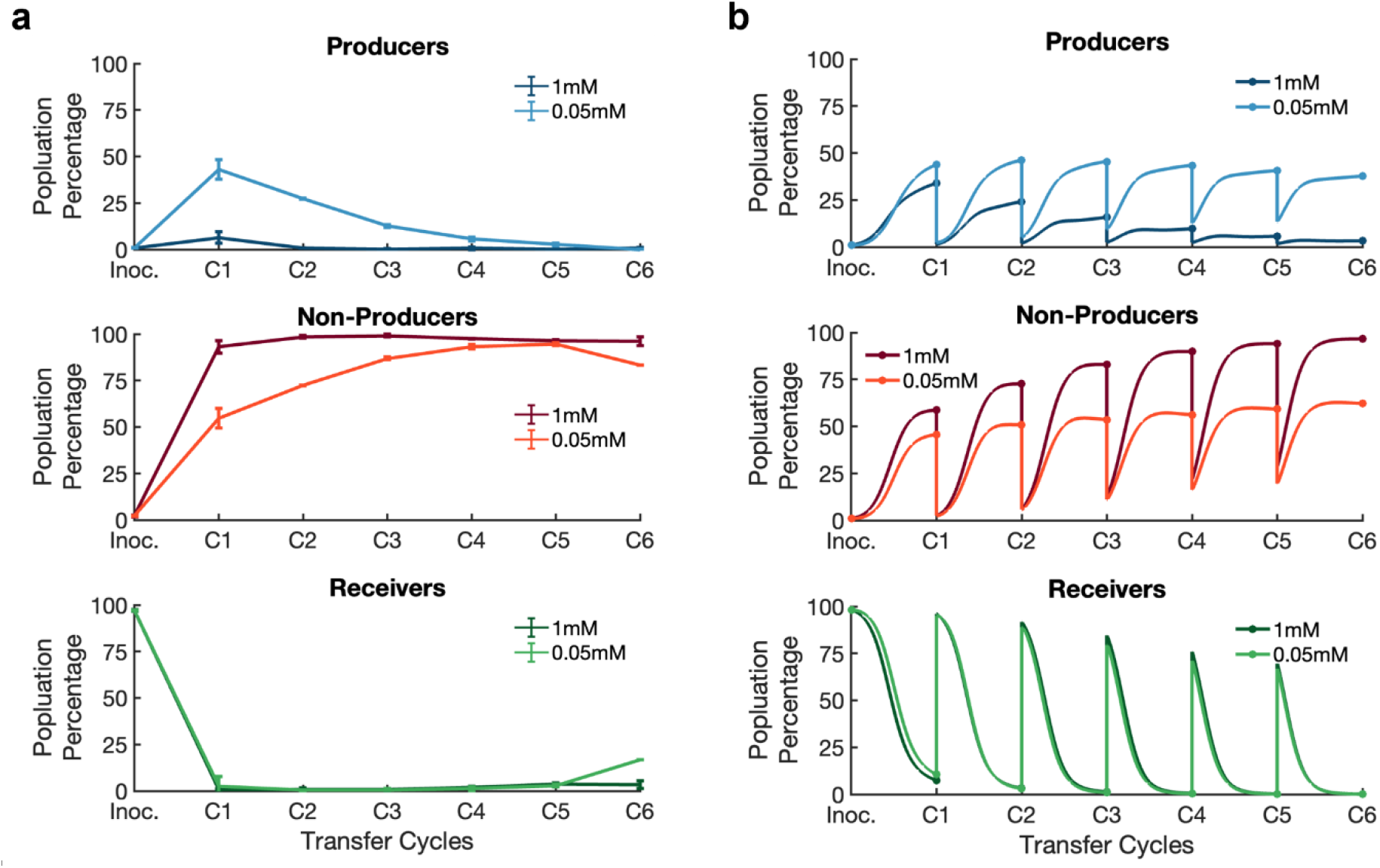
A strategy to enrich for Producers by lowering IPTG concentration to increase competition from Receivers. **a** The cell counts corresponding to the Producer, the Non-Producer, and the Receiver population percentages are used to compare the growth trends of the two culturing conditions. After 12 h of growth, the cell counts for each population were measured for 6 cycles. At the end of the first growth cycle, the Producer cell counts at 0.05 mM IPTG induction condition is ~7-fold higher than at 1 mM IPTG condition (from about 6.3% to 42%). As we iterate the growth cycle, Producer percentage drops in both conditions. By the end of the 6^th^ cycle, with Producers mostly gone, the Receivers percentage starts to rise, which shows the competition between the Non-Producers and Receivers. (n = 2, error bars = s.d.) **b** Simulation of equations (S6) for the three populations at two different IPTG concentrations. Parameters used are listed in Table S2.

As shown by the Receiver percentages at the end of cycles 1 to 5, the fast effective diffusivity and the accumulation of AHL molecules still leads to the elimination of Receivers, even at the lower IPTG concentration. To further lower AHL accumulation in the system, we diluted the Producer population size by 10-fold in the initial co-culture, with a 0.1:1:100 starting ratio of P to N to R. The results showed that most of the Receivers survived with the lower Producer population and further limited Non-Producer growth **(Fig. S13)**. However, the Producers only grew to ~3% of the population at the end of the first cycle. When iterating the growth cycle with spiking in of a fixed number of Receivers, we observe that both Producer and Non-Producers are being overtaken by the Receivers, which is presumably due to the increased competition of the Receivers added at the end of each cycle. Our results suggest that, without a more sophisticated adaptive control on the concentration of Receiver spiked-in, the competition-based spatial segregation produced through the microdroplet system alone is not sufficient to enrich for slow-growing rare Producers, where the public good diffuses fast and degrades slowly.

### A fluorescent-activated microdroplet sorting strategy to continue enriching the Producer subpopulation

For a system where the public good diffuses extremely fast in the timeframe of cell growth, we incorporated a fluorescence-based sorting module into the competition-based microdroplet system. The fluorescence sorting was used to screen for microdroplets with green fluorescence (Receiver cells) **(Fig. S14)**. As before, we first co-encapsulated the Producers and Non-Producers at low density (*λ* ≪ 1) with Receivers at higher density (*λ* > 1) to achieve spatial segregation of the Producers from Non-Producers while keeping the Receivers present in most droplets **(Fig. 5a)**. After incubation, the P+R microdroplets would have fewer Receivers than the ones from N+R microdroplets, assuming some of the AHL is privatized because of the sequestration by Receivers and the extremely rare Producers.

**Figure 5:**
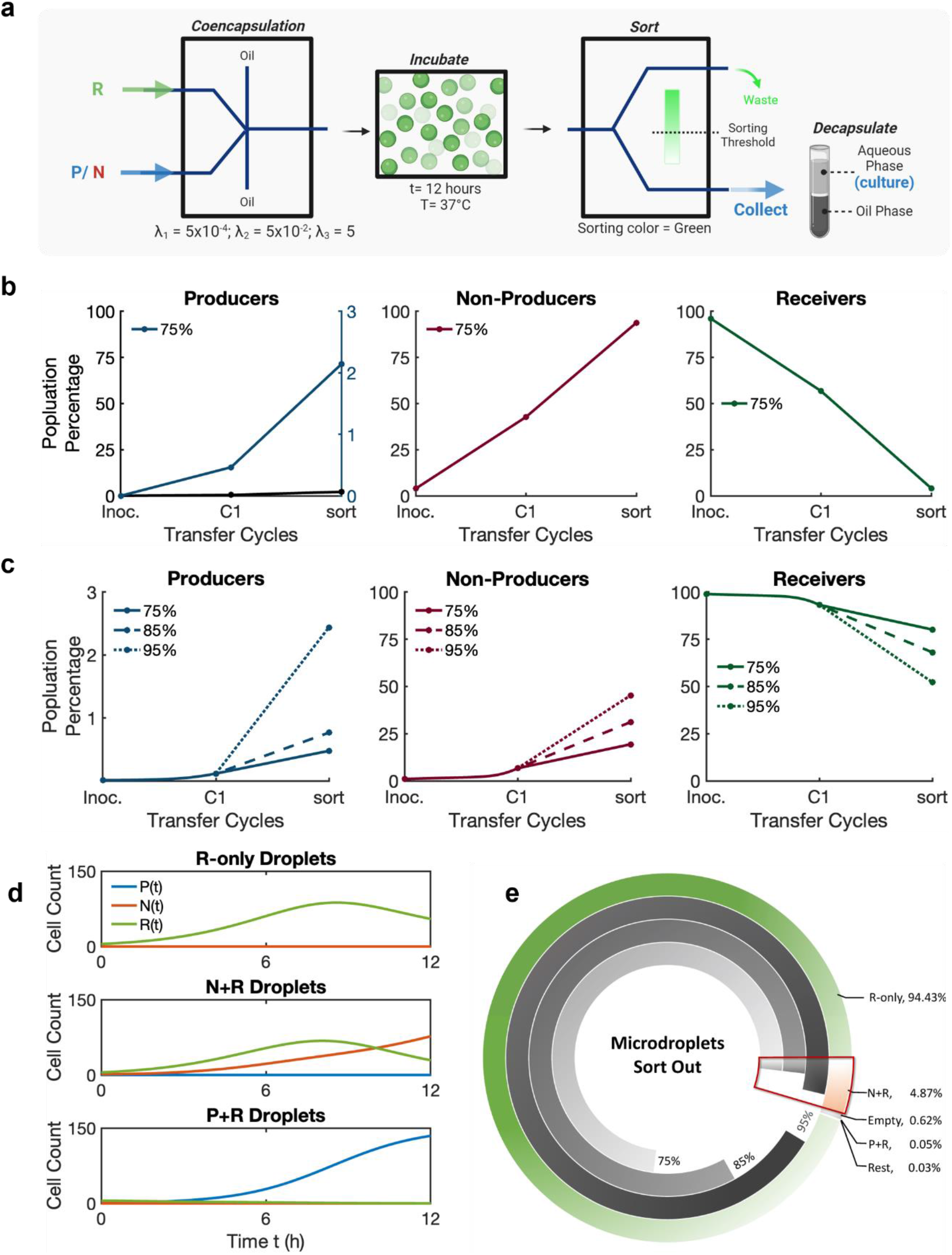
Enriching for rare Producer population by incorporating Fluorescence Activated Droplet Sorting (FADS) into the competition-based experimental strategy. **a** Schematic of the microfluidic platform including microdroplet generator for co-encapsulation and FADS. **b** Population percentages for the Producers, Non-Producers, and Receivers at inoculation (Inoc.), 12 h of incubation (C1) followed by the fluorescent sorting (sort) at 75% thresholds. Total number of droplets screened, n= ~380k microdroplets. **c** Simulation of equations (S6) and (S7) with three different sorting thresholds: 75%, 85%, and 95%. **d** Simulation result for population counts of each strain from all droplet that contains: R-only, N+R and P+R. This shows that under the middle diffusion scenario (β = 1/2), Receivers are effectively competing with Non-Producers but not Producers. **e** Demonstration of the composition of the droplets that get sorted out at different thresholds. The most-outer ring describes the statistics of microdroplet by catoegory after initial incubation for λ_1_ = 5 × 10^−4^ for Producers, λ_2_ = 5 × 10^−2^ Non-Producer, and λ_3_ = 5 for Receivers. The three inner rings show the portion of the droplets from each category that is sorted out (grey-scale colored) and stays in (white colored) for 75%, 85%, and 95% (inner to outer) thresholds. See Table S3 for detailed simulation results.

Here, the initial population densities were chosen such that *λ*_1_ = 5× 10^−4^ for Producers, *λ*_2_ = 5 × 10^−2^ for Non-Producers, and *λ*_3_ = 5 for Receivers, (a 0.01:1:100 starting ratio of P to N to R). The cells were inoculated and grown in microdroplets identical to the previous experiments. After incubation, the microdroplets are injected into a tapered channel in the sorting module **(Fig S15)**, where the voltages corresponding to the green-fluorescent microdroplets are recorded **(Fig. S16)**. Based on the recorded voltage distribution, we can set cutoff voltages to sort fluorescent microdroplets above the cutoff value **(Fig. S17)**. The stringency of sorting can therefore be controlled by setting the voltage cutoff from the detected fluorescent signals. A sorting threshold of 75% means that three-quarters of the highly fluorescent green microdroplets detected were sorted out of the system. The remaining 25% of the microdroplets with lower green fluorescence are collected for the cell count. After one round of sorting at 75% thresholds, the experimental data shows that the Producer population increased 50-fold, from 0.4% to over 2% **(Fig. 5b)**.

When modeling the system with the low initial Producer counts (*λ*_1_ = 5 × 10^−4^), sequestration of signals takes a relatively larger effect on the effective diffusion, leading to a significantly lower percentage of signals leaving the microdroplet. In particular, we consider a middle case scenario described by equation (3), with *β* = 1/2. That is, we assume half of the signals won’t leave the microdroplet where they are produced, while the other half will be shared immediately with all the microdroplets. Under this scenario, Receivers are effectively competing with Non-Producers but not Producers, as shown in **Fig. 5d**. This simulation results also suggest that R cells are higher in the R-only and N+R droplets than the P+R droplets. After sorting out the droplets with higher R cells, the Producer percentage increases **(Fig. 5c)**. Meanwhile, when we increase the sorting threshold further, the sorting is more efficient as we begin to sort out the N+R microdroplets as shown in **Fig. 5c, e**. However, experimentally increasing the thresholds can be difficult currently as there are tradeoffs and technical limitations in detecting fluorescence and errors in selecting microdroplets of interest at fast flow rates **(Fig. S18)**. For example, a very rare event such as a positive occurring at a frequency of 1 in a million in an emulsion of 1 million microdroplets may not be efficiently isolated at high sorting speeds and slower sorting speeds will dramatically increases sorting times. While spatial segregation afforded by microdroplets provide a window of opportunity for Producer enrichment, incorporating sorting module extends the opportunity further by eliminating the microdroplets with Receivers and Non-Producers. Consequently, we can increase the Producer percentage beyond the benefits afforded by growing the populations in structured microdroplets.

## Conclusion

Understanding how natural and synthetic microbial communities arise and how to program them for specific tasks would accelerate innovations in biotechnology, medicine, and biomanufacturing. Spatial structure is an important aspect of natural multi-strain communities and contributes strongly to their robustness and diverse phenotypes^35–41^. Earlier studies have shown that when secreted signals or cell types are highly constrained (low or no diffusion) within microdroplets, slow-growing but desirable high-yielding cells producing valuable biomolecules can be enriched efficiently^23,34^. However, it is likely that many small molecules of interest, especially signaling molecules or antimicrobials could readily diffuse between microdroplets and alter population dynamics. To engineer both natural and synthetic microbial communities will require an understanding of the molecular principles of social interactions as well as strategies to shape communities to desired characteristics. We used a three-member microbial synthetic community in which the Producers secrete AHL as signaling molecules that can diffuse between the microdroplets to investigate the role of signaling within a defined synthetic community and develop experimental and mathematical modeling approaches to control emergent Producer population dynamics.

Using emulsions of microdroplets as individually compartmentalized microenvironments, we were able to quantitatively assess and model the interplay of population dynamics when diffusible signals are present. Of particular importance was the development of strategies that could allow for the enrichment of valuable Producer subpopulations despite moderate to fast diffusion. The production of valuable biomolecules or controlling phenotypes of synthetic communities for translation applications like bioremediation will typically incur fitness costs to the Producer that invite the rise of cheating Non-Producers. We showed that during serial propagation in a spatially structured environment, the window of opportunity for the selection of the AHL Producer population is small as the Non-Producers can rapidly outcompete Producers. This is caused by rapid diffusion of AHLs, which, in turn, leads to reduced Receiver competition in all the microdroplets. While AHL production is metabolically costly to the Producers, the benefits of reduced Receiver competition are shared by both the Producers and the Non-Producers alike. In the absence of spatial structure, as in the case of a well-mixed batch culture, Non-Producers rapidly outcompete both Producers and Receivers. Even with fast diffusion, the spatial structure imposed by microdroplets did slow the rise of Non-Producers. Unlike in batch culture, the Non-Producers within microdroplets are limited by the physical confinement of the microdroplet and cannot access the entire culture volume of the emulsion and therefore expand more slowly.

The mathematical model developed using this microbial synthetic system and our experimental findings showed that Producer success in the community when AHL diffuses among the microdroplets is transient. This may seem counterintuitive as it might have been expected that the community might reach a stable steady-state. Our data clearly show that in the case of rapid diffusion, cheaters (Non-Producers) will eventually force the Producers and Receivers to extinction. Importantly, the simulation also suggested that the window of opportunity for Producer enrichment and isolation can be affected by different environmental factors such as initial population size, AHL secretion rate, sensitivity of the population to the signal, composition of the droplets, the duration of the interaction, and dilution factor. Our results pointed toward several parameters that could be adjusted to increase the probability of extending the window of opportunity for the AHL Producer to succeed.

We tested predictions of the simulation and showed that we could extend the success of the Producers by reducing AHL production, and by using Fluorescence Activated Droplets Sorting (FADS) sorting. Using FADS to sort for microdroplets with lower green fluorescence (Receivers), we observed a 50-fold improvement for the enrichment of Producers. We deliberately chose to sort the microdroplets based on Receiver fluorescence (indirect) rather than the Producers (direct, the cell type of interest) with a specific application in mind. Translational applications of synthetic communities to produce valuable molecules will likely make use of the abundant diversity of both natural and engineered microbial life. Indirectly enriching for Producers within the context of fast diffusion could be used to enrich for wild strains of bacteria without the need for genetic manipulation such as introducing a fluorescent reporter. By avoiding genetic manipulation of the Producer or strain of interest, we can dramatically increase our ability to evaluate wild strains and communities rapidly and at scale. Further, if we define a Producer as a strain, strain variant, or community that produces a valuable activity or molecule(s), then using an indirect approach to enrich Producers allows an investigator to engineer a reporter strain or molecule that reports on the desired activity. In experiments where the population of interest is labeled, it can be sorted directly^33,42–44^, and therefore population ratio increases dramatically. By evaluating an indirect method, we develop opportunities to screen for bacteria of interest based on their activity towards a directed task.

Given sufficient accuracy of the sorter, a better enrichment of Producers can be achieved through increasing the sorting threshold and/or iterating the growth cycles with sorting. Since the indirect sorting is done based on the survived Receiver’s fluorescence in droplets, the upper limit of the sorter for enrichment is bounded by the number of Producers that’s high enough to kill all Receivers in the system. Therefore, when enriching for Producers with iteration, one can increase the Receiver cells spiked-in to further extend the enrichment window for Producers. With accurate sorting and adaptive Receiver spike-in, the platform is theoretically capable of enriching for 1 Producer from 1 million Non-Producer cells.

In summary, we have tested and validated that enrichment of rare but potentially valuable subpopulations within a much large population of competitors and cheaters can be achieved even when the desired rare subpopulation is not directly labeled. As microfluidic technologies for microdroplet production, fluorescence sorting, and microdroplet capture improve, new opportunities for selection and enrichment of rare strains and synthetic microbial communities with desirable phenotypes will also improve.

## Methods

### Bacterial strains and growth conditions

All experiments were conducted in strain JS006-ALT (parent strain: *E. coli* K-12 BW25113 Δ*lacI* Δ*araC* (*tetR*^−^) with *lacI*, *araC*, and *tetR* integrated under the control of constitutive promoters^45^. The plasmids were constructed using either golden gate or Gibson assembly. Plasmids were transformed into JS006.ALT cells using heat shock transformation protocol using chemically competent cells. Transformants were plated on agar plates with appropriate antibiotics. The antibiotic concentration used for all experiments are as follows: ampicillin 100 μg/mL, kanamycin 50 μg/mL. Transformation plates were incubated at 37 °C for 16 - 18 h. Colonies were picked from the plate and inoculated in 5 ml of LB media for overnight cultures. All the plasmids and the corresponding strains used in this study are listed in Table 1.

**Table 1:** Plasmids used to construct the synthetic community.

| Strain | Plasmid | Gene Cassette | Resistance | Ori |
| --- | --- | --- | --- | --- |
| Producer - Plasmid 1 | C162b | P<sub>lac</sub>-*lasI*-LAA | kan<sup>R</sup> | pMB1 |
| Producer - Plasmid 2 | RA016 | P<sub>lq</sub>-*sfCFP* | amp<sup>R</sup> | p15A |
| Non-Producer - Plasmid 1 | RG01 | P<sub>lac</sub>-*lasI*<sup>trunc</sup>-LAA | kan<sup>R</sup> | pMB1 |
| Non-Producer - Plasmid 2 | RG02 | P<sub>lq</sub>-*mCherry* | amp<sup>R</sup> | p15A |
| Receiver - Plasmid 1 | C158 | P<sub>las-lac</sub>-*holin*; P<sub>las-lac</sub>-*lysin*-LAA | kan<sup>R</sup> | pMB1 |
| Receiver - Plasmid 2 | RG03 | P<sub>lq</sub>-*sfGFP*; P<sub>lq</sub>-*lasR* | amp<sup>R</sup> | p15A |

The Producer contains a plasmid, C162b coding for IPTG induced expression (P_lac_) of quorum sensing (QS) gene *lasI* and a second plasmid, RA016 coding for the constitutive expression (P_Iq_) of *sfCFP* for strain identification. *lasI* gene expression is driven by a LacI promoter with the modified bicistronic design ribosome binding site^46^. LasI synthases produce *N*-(3-Oxododecanoyl)-L-homoserine lactone (C12-oxo-HSL) molecules that can diffuse out of the cell into the media and nearby cells. The *lasI* gene was tagged with the original LAA ssrA degradation tag and had the iGEM registry B0014 terminator ^47^. This plasmid also contained a kanamycin resistance gene for selection, pMB1 origin, and ROP element that reduces the copy number^48^. The Receiver strain contains a plasmid, RG03 coding for constitutive expression (P_Iq_) of the C12-HSL response regulator, *lasR*, and *sfGFP* for strain identification. It also has a second plasmid, C158 coding for two toxic genes, *holin* and *lysin* inducible by IPTG and C12-oxo-HSL (P_las-lac_). Expression of these genes in the presence of AHL molecules kills the Receivers. The *holin* and *lysin* genes were tagged with the original LAA ssrA degradation tag and had the iGEM registry B0014 terminator^47^. This plasmid also contained a kanamycin resistance gene for selection, pMB1 origin, and ROP element that reduces the copy number. The Non-Producer strain has a plasmid, RG01 coding for IPTG induced expression (P_lac_) of a disrupted *lasI* gene which prevents active AHL production and therefore cannot kill the Receivers. The fourth amino acid in the *lasI* gene, glutamine (CAA) was modified into a stop codon (TAA), resulting in the early termination of AHL synthase. The Non-Producers have a constitutive P_Iq_ promoter and modified bicistronic design ribosome binding site that drives the expression of *mCherry* for identification^46^. This plasmid, RG02 also contained a kanamycin resistance gene for selection, pMB1 origin, and ROP element that reduces the copy number^48^.

All *E. coli* strains were grown in LB media containing kanamycin (50 μg/ml) and ampicillin (100 μg/ml) at 37 °C in triplicates. The next day, the cultures were then diluted to OD 0.05 before further diluting 100-fold into the fresh medium in 96-well plates. For the co-culture experiments, appropriate volumes of the above culture were mixed before loading on a 96-well plate. The optical densities were monitored using a microplate reader (Tecan Spark) at 37 °C for 24 h, with data recorded every 10 minutes. The final graph of OD or fluorescence vs time was plotted using the average of the 3 biological replicates with standard deviation (s.d.). Doubling times were calculated from the growth curve between the OD 600 interval of 0.2 and 0.4.

### Fabrication of microfluidic devices for microdroplet production and sorting

We fabricated a microfluidic microdroplet generator and a sorting device **(Fig. S13)** in PDMS (poly(dimethylsiloxane)) following conventional soft lithography techniques using the designs from Mazutis et al.^33,50^. Briefly, the SU8-2025 photoresist (Kayaku Advanced Materials (formerly MicroChem Corp.) was spin-coated onto silicon wafers and patterned by UV exposure through a photolithography mask and subsequent development (SU-8 developer; MicroChem Corp.). From the photoresist masters, the microfluidic devices are made using a translucent, gas permeable PDMS (Sylgard 184, Dow Corning) at a 10:1 (w/w) polymer to crosslinker ratio. PDMS is degassed, then poured onto the master, and cured at 80 °C for 16 hours. Afterward, the cured PDMS is peeled off of the master. Access holes for tubing on the PDMS monoliths were punched using a Harris Unicorn biopsy punch (inlet ports - 0.75 mm, outlet ports - 1.5 mm, and electrode ports – 0.5 mm). The PDMS monolith was then bound to 1.5 cover-glass by activating both surfaces with oxygen plasma cleaner (Harrick Plasma, PDC-001). The channels are made hydrophobic by using Aquapel to treat the surfaces of the channel. Then, the channels were immediately rinsed using Novec HFE-7500 (3M) to remove unreacted Aquapel. For the microdroplet sorting device, indium tin oxide (ITO) coated glass slides were used (Delta Technologies). The electrodes for this device were fabricated by inserting a low-melting soldering wire into the designed channels.

### Encapsulation of cells into microdroplets

The microfluidic microdroplet generator has an inlet for the oil phase and two inlets for the two different bacterial cultures that are targeted to the flow-focusing junction to form uniform microdroplets. With the aqueous flow rates of 10 μl/min and oil flow rates of 15μl/min, we can produce the desired microdroplets of 40 ± 2 μm in diameter (n = 1108, volume = ~33.5 pL) at the speed of ~5000 Hz. A movie made to illustrate droplet production is provided in Supporting Information, **Video S1**. Bacterial culture was used as the aqueous phase whereas Novec HFE 7500 fluorinated oil (3 M, Maplewood, MN, USA) with 1.5% PicoSurf surfactant (Sphere Fluidics, Cambridge, UK) was used as the oil phase. The microdroplets generated were collected in 50 ml tubes through an outlet on the microfluidic device. The microdroplets are then sampled and imaged using a VitroCom glass capillary tube (Mountain Lakes) under a 20X objective on a Ti2E inverted microscope (Nikon). When samples from the microdroplet collection tubes are visualized at different time points in a capillary tube, the location of the microdroplets to each other is not maintained in this process. Fluorescence was imaged using either EGFP: Excitation: 480/30 nm, Emission: 535/20 nm; mCherry: Excitation:560/40 nm, Emission: 635/60 nm; or ECFP: Excitation: 435/20 nm, Emission: 480/30 nm filters (Chroma).

### Microdroplet sorting

A microfluidic sorting device has an inlet for reinjection of the formed microdroplets and an inlet for the fluorinated oil that separates microdroplets for efficient sorting. To achieve this, the microdroplets were injected at a flow rate of 20 μl/h and the oil phase was injected at the rate of 480 μl/h. The spaced microdroplets from the pico-injection module **(Video S2)** enter the sorting module, where the microdroplets can be sorted into a collection or a waste chamber via di-electrophoresis using a custom LabVIEW program developed by Mazutis et al. that records microdroplet fluorescence and trigger the electrodes during sorting^33^ **(Fig. S14, S15)**. A movie made to illustrate microdroplet sorting is provided in the Supporting Information **(Video S3)**. The data acquisition rate for this system is 100 kHz. The throughput for microdroplet sorting was ~100 microdroplets/sec. The voltage thresholds were decided based on signal profiles from positive and negative control microdroplets, and sorting efficiency was verified by sorting bright microdroplets. The electrodes fabricated within the microfluidic device induce a non-uniform electric field that creates a dipole effect in the microdroplet. If a microdroplet carrying a detectable fluorescent cell passes through a point in the channel irradiated by a corresponding wavelength laser, the electric field is turned on. The sorting junction is designed such that the collection chamber has higher resistance than the waste chamber^33^. Therefore, in the absence of an electric field, all the microdroplets enter the waste chamber. A digital CMOS camera and a high-speed camera (Phantom V7.2) were used to capture images during microdroplet formation and microdroplet sorting. The high-speed camera was connected to the LabVIEW software and vision module.

### Serial Propagation in emulsion microdroplets

After overnight growth, the cultures were washed and inoculated in fresh LB media with relevant antibiotics to reach a McFarland unit of 5.0 for Receivers and 0.1 for Producers and Non-Producers. Initial population ratios were determined by plating dilutions on LB agar plates. Based on the measured concentrations, the pre-cultures were diluted to 1.5*10^8^ cells/ml of Receivers, 1.5*10^6^ cells/ml of Producers, and Non-Producers each. This corresponds to Receivers *λ*_3_ = 5, Producers *λ*_1_ = 0.05, and Non-Producers *λ*_2_ = 0.05. About 500 μl of microdroplet corresponding to 15 million microdroplets with an average diameter of ~40 μm was generated. All the experiments in microdroplets were replicated in duplicates. The microdroplet cultures were incubated without shaking to avoid droplet coalescence. After overnight growth, the aqueous phase containing the cells was separated from the oil phase by adding 300 μl of perfluoro-octanol. The mixture was gently mixed by flicking the tube, then allowed to settle for 5 min. The top aqueous layer was collected and used for further analysis, as inoculum for the next propagation, and for making a glycerol stock solution (supplemented with 30% v/v glycerol, stored at −80 °C). The cultures were washed with fresh media, diluted 50 folds, and spiked with Receivers between each transfer. Microscopic images and cell counts were recorded at the end of each cycle.

### Serial Propagation in suspension

Serial transfers in the test tube were performed in a similar fashion as the emulsion microdroplet experiments. All the test tube experiments were replicated in triplicates. After overnight growth, each strain in monoculture was inoculated in 5 ml of fresh media to reach a McFarland unit of 5.0 for Receivers and 0.1 for Producers and Non-Producers. These concentrations mimic the initial concentrations in the inoculated water-in-oil microdroplets. Suspension cultures were incubated with shaking at 200 rpm and 37 °C to ensure well-mixed conditions. After growth, the suspension cultures were used for further analysis, as inoculum for the next propagation, and for making a glycerol stock solution (supplemented with 30% v/v glycerol, stored at −80 °C). The cultures were washed with fresh media, diluted 50 folds, and spiked with Receivers between each transfer. Microscopic images and cell counts were recorded at the end of each cycle.

### Bacterial cell count using agarose pads

We used agarose pads to immobilize the cells, allowing us to perform multi-color fluorescence microscopy imaging. We followed the protocol for making the agarose pads from Skinner et al.^51^. Briefly, 1% agarose in 10 ml PBS was microwaved for about 30 seconds, until the agarose was completely dissolved. The agar solution was pipetted onto a chamber prepared using stacking glass slides. After 45 minutes at room temperature, the agarose solidifies, which is then cut into smaller squares. For imaging, the bacterial cells, 1 μl of culture was pipetted onto a glass coverslip and covered by an agar pad. These agar pads were then imaged on an inverted fluorescence microscope; taking multiple images in different areas and counting at least 500 cells to get a better representation of the data. Fluorescence was imaged using either EGFP: Excitation: 480/30 nm, Emission: 535/20 nm; mCherry: Excitation: 560/40 nm, Emission: 635/60 nm; or ECFP: Excitation: 435/20 nm, Emission: 480/30 nm filters (Chroma). Since the signals from the EGFP channel can bleed through to the ECFP channel, the exposure time and the LUIs were carefully selected such that true green-fluorescent and cyan-fluorescent signals are counted. The cell count was analyzed using the Nikon Elements software.

### Image analysis

For each position, images in bright-field and fluorescence using either EGFP, mCherry, or ECFP fluorescent channels while microdroplets were immobile inside rectangular capillary tubes (Vitrocom, USA). Images were recorded with a PCO.edge 5.5m camera (PCO, Germany). Images were analyzed for microdroplet size using custom python code. Microdroplets were identified from the bright-field images using the Hough transformation algorithm (OpenCV 3)^35^.

### Statistical analysis

Replicate numbers of the experiments (*n*) are indicated in the description of the method. Data in the figures are presented as mean ± s.d. Fluorescence microscopy images are representatives of the images from multiple experimental replicates. All statistical analysis was performed using Graphpad Prism and MATLAB.

## Supporting information

Video S1 Microdroplet generation

Video S2 Picoinjection

Video S3 Microdroplet Sorting

## Abbreviations

AHL: Acyl Homoserine lactone
QS molecules: Quorum Sensing molecules
HSL: Homoserine lactone
P: Producer
N: Non-Producer
R: Receiver
FADS: Fluorescence Activated Droplet Sorting
IPTG: isopropyl β-d-1-thiogalactopyranoside
C12-oxo-HSL: *N*-(3-Oxododecanoyl)-L-homoserine lactone)
PDMS: poly(dimethylsiloxane)
*sfCFP*: super-folder Cyan Fluorescent Protein
*sfGFP*: super-folder Green Fluorescent Protein
s.d.: Standard Deviation
Inoc.: inoculation

## Author Contributions

Y.S. and R.G.P. conceived of the project. R.G.P., Y.S., and G.F. designed the experiments. G.F. constructed and simulated the mathematical model. R.G.P. and G.F. analyzed the data. R.A. and A.H. consulted on microfluidics and the optical instrumentation setup. R.G.P. performed all the experiments. R.G.P., G.F., and Y.S. wrote the original manuscript. All authors reviewed and commented on the manuscript.

## Competing Interest

The authors declare no competing financial interests.

## Supporting document information

The supporting information is available free of charge on the ACS Publications website at DOI:

Supplementary Figures S1-S18, Supplementary Table 1-3, and Mathematical model supplement (PDF)
Video S1 (Microdroplet production)
Video S2 (Microdroplet pico-injection)
Video S3 (Microdroplet sorting)

## Data availability

The authors declare that the data underlying the findings of this study are available within the paper and its Supplementary information files and are available upon request.

## Code availability

All code used to create the figures is available at https://github.com/BridgetFan/Enrichment.git.

## Acknowledgments

The authors also acknowledge the use of resources of the Shared Equipment Authority at Rice University for this work. This work was supported by NIAID Award R01A1080714 (Y.S.), NSF award 1936770 (G.F.), DMS-1662290 (M.R.B) and NIH grant and R01GM144959 (M.R.B). The figures and schematics were created with MATLAB, GraphPad, BioRender.com, and Excel.

## Notes

### Competing Interest Statement

The authors have declared no competing interest.

