## Supplemental Information for "Indirect enrichment of desirable, but less fit phenotypes, from a synthetic microbial community using microdroplet confinement"

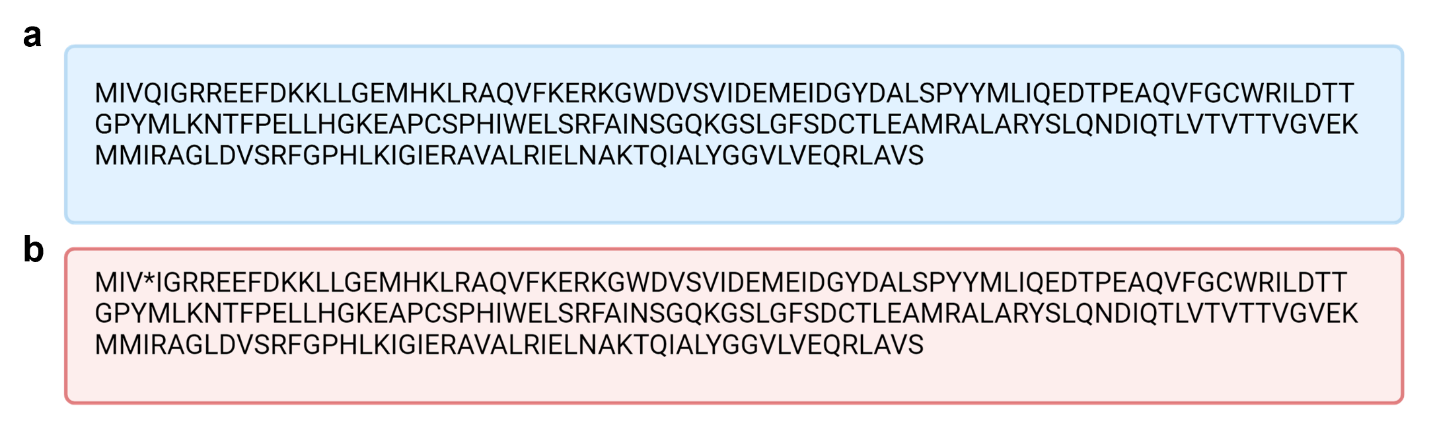

**Figure S1:** *Amino acid sequence of lasI gene in Producer and Non-Producers.* **a** Amino acid sequence of *lasI* gene in Producer strains, Plasmid C162b. **b** Amino acid sequence of *lasI* gene in Non-Producer strains, Plasmid RG01. The fourth codon of the LasI, glutamine (CAA) was modified into a stop codon (TAA), resulting in the early termination of AHL synthase synthesis.

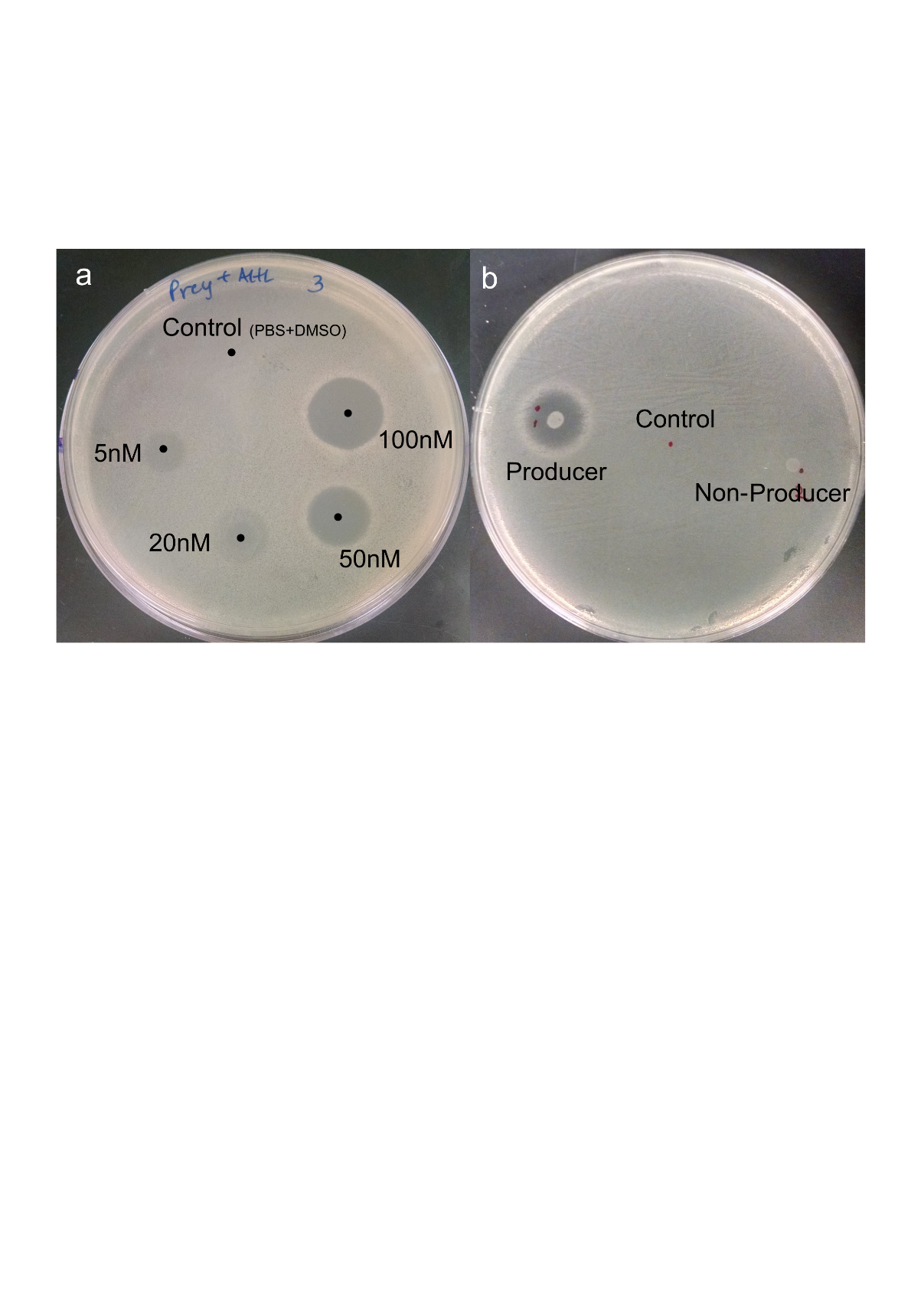

**Figure S2:** *Zone of inhibition (ZOI) assay on agar plates overlaid with the Receiver cells.* **a** The left plate shows the ZOI as the Receiver strain responds to different concentrations of AHL molecules. **b** The right plate shows the response of the Receivers to the Producers and the Non-Producers respectively. When the Producer cells were spotted on an agar plate overlaid with the Receiver cells, we see a clear zone of inhibition (ZOI) or killing, whereas the Non-Producer cells do not induce Receiver cell lysis as the AHL production is disrupted.

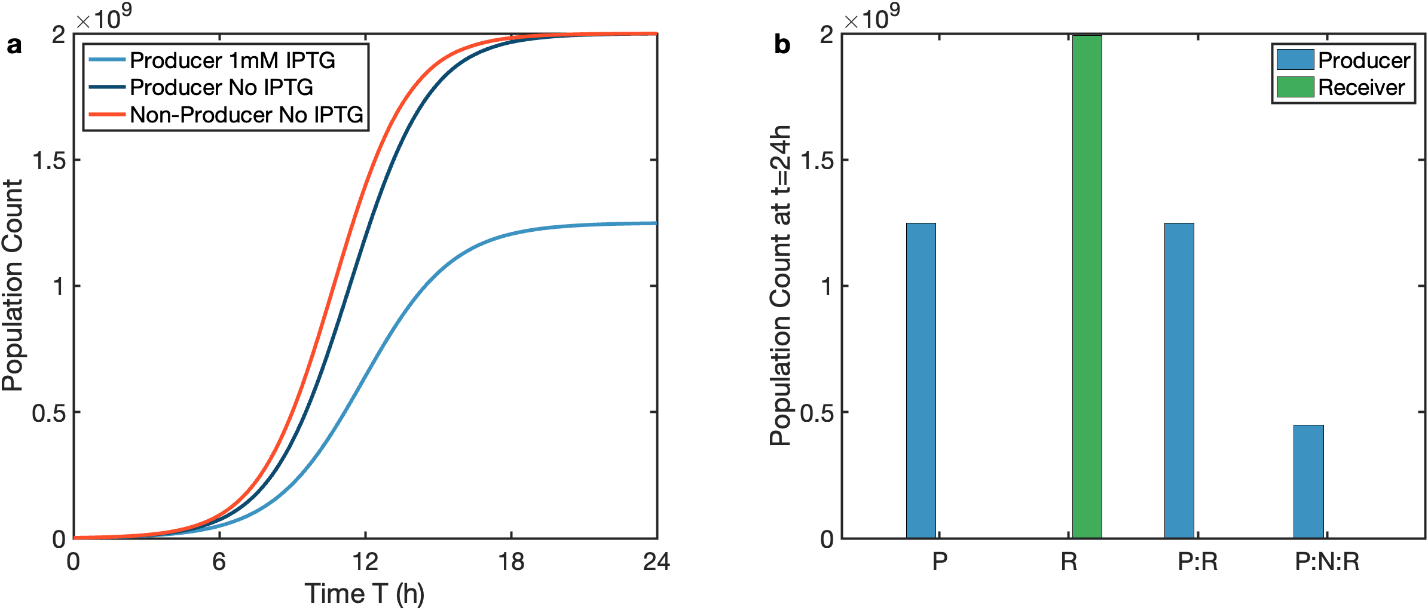

**Figure S3:** *Simulation result of equations (S1) for the growth of Producers in different conditions.* **a** Simulation result for Producers and Non-Producers under different IPTG conditions. **b** Simulation result of the ending population count at t = 24 h for Producers (P) and Receivers (R) with either monoculture or co-culture P:R and P:N:R under 1 mM IPTG inducer level. Parameters used are listed in Table S2.

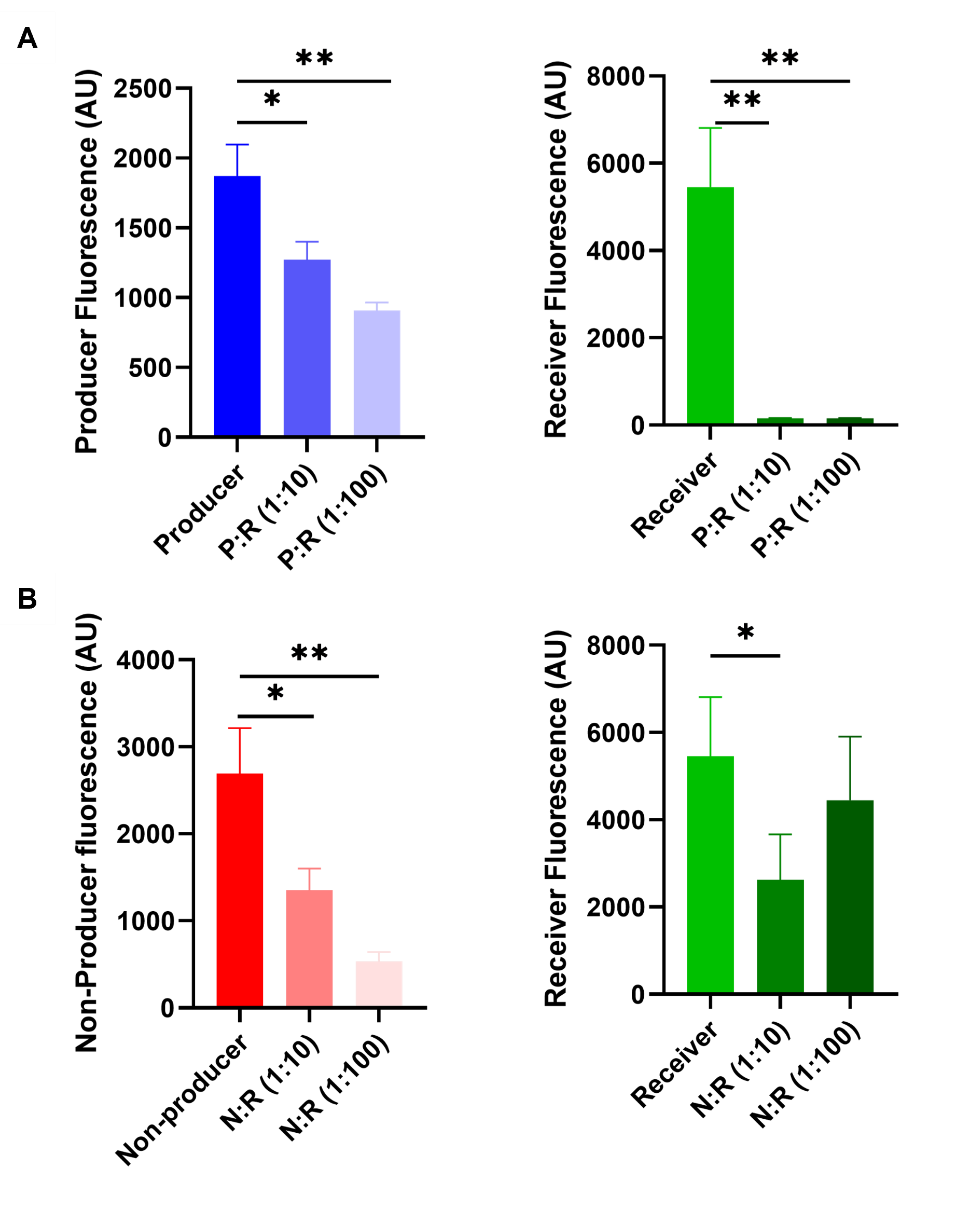
**Figure S****4:** *Producers can kill up to 100-fold excess of Receivers.* The graphs show comparisons of the final population density of the Receivers in co-culture with Producers (P:R), measured using their respective fluorescence proteins. At 1 mM IPTG induction, when Receivers are co-cultured at 10-fold or 100-fold excess of Producers (P:R), all the Receivers are killed (Receiver fluorescence is close to the background). n = 3, error bars = s.d., Unpaired t test, *P* value * .016, *P* value **.002

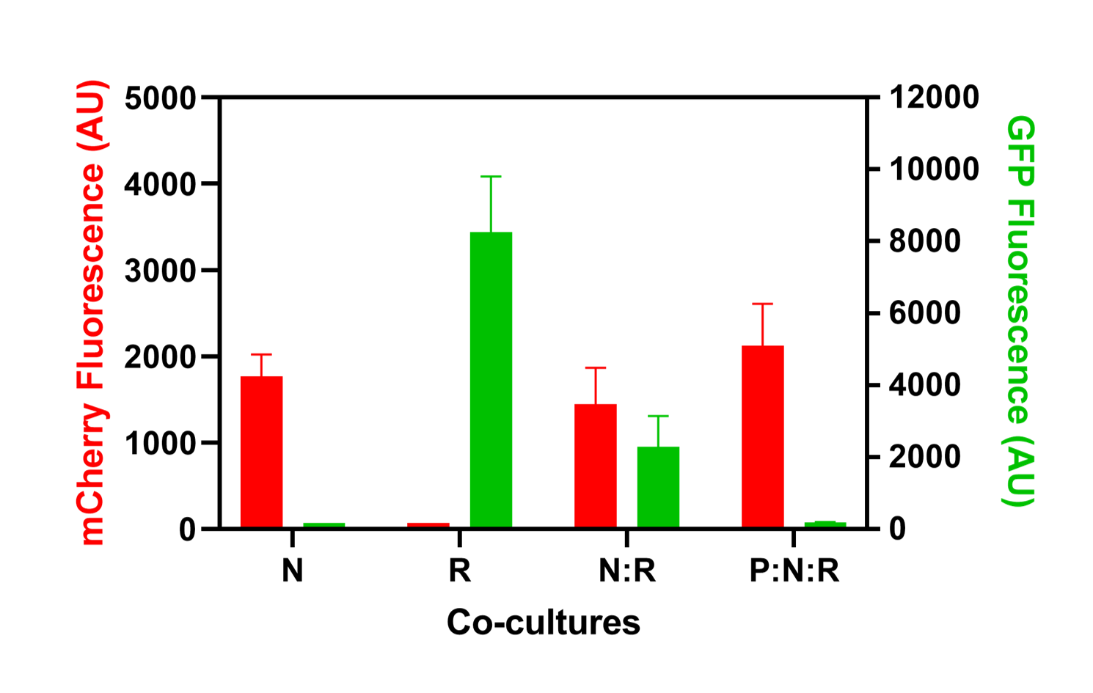

**Figure S5:** *Non-Producers and Receivers compete for common resources.* Comparisons of the final population density of the Non-Producers in co-culture with Receivers and Producers, measured using their respective fluorescence proteins. N is the Non-Producer grown in mono-culture at 1mM IPTG after incubation at 37^o^C for 16 -18 hours. R is the population density of Receivers after incubation under identical conditions. N:R is the Non-Producer-Receiver co-cultures, and P:N:R is the Producer-Non-Producer-Receiver cocultures incubated at identical growth conditions with an initially equal population ratio.

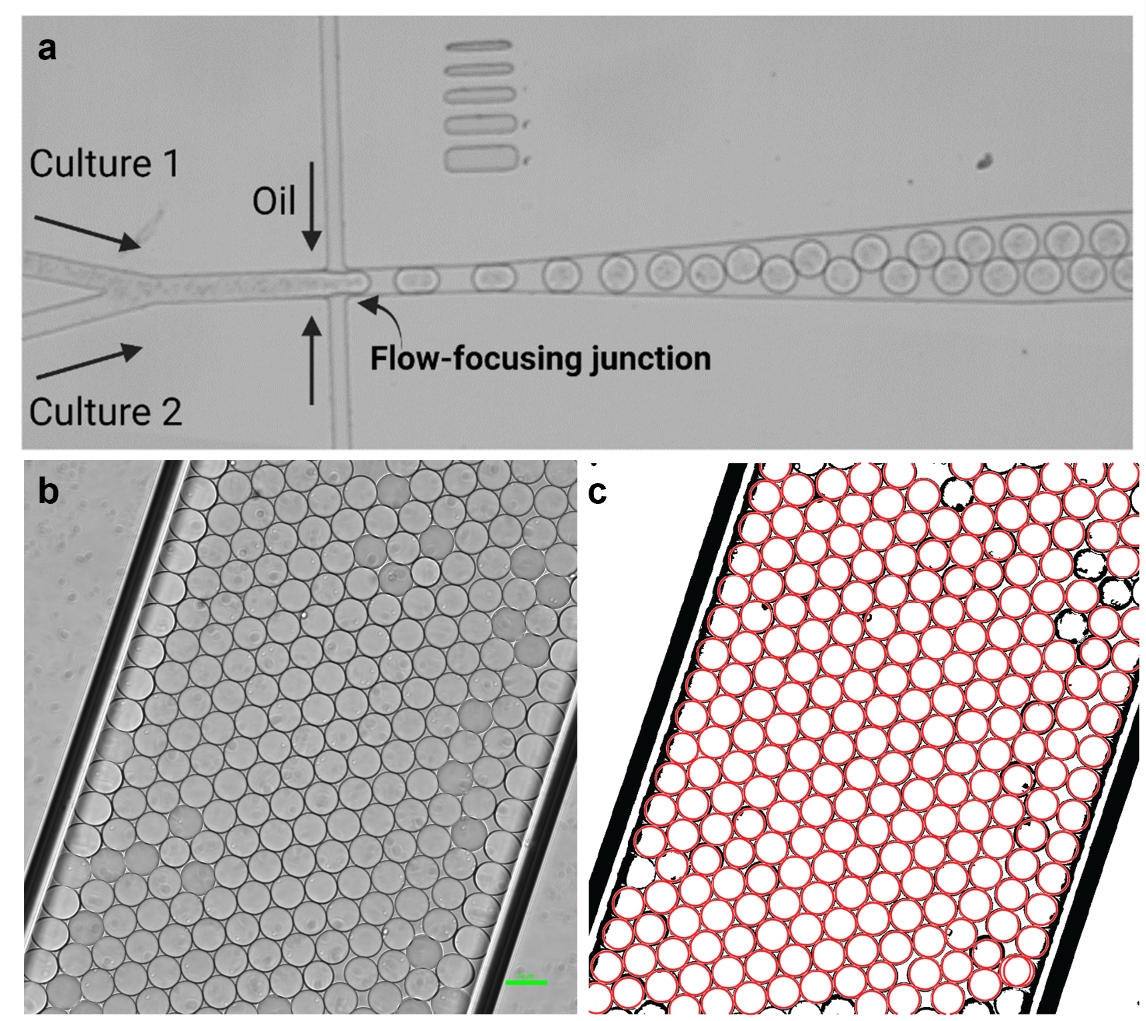

**Figure S6:** *Microfluidic droplet generator device.* **a** Flow-focusing junction where the oil and the culture phases are focused to form droplets of about 40 μm diameter. Visualized using brightfield microscope (20x). Scale bar: 50 µm. Brightfield image of microemulsion droplets **(a)** and image analyzed **(b)** for automated droplet identification and diameter measurements using a custom code in MATLAB. Droplets were identified from the phase contrast image using a Hough transformation algorithm (OpenCV 3). n = 235 microdroplets.

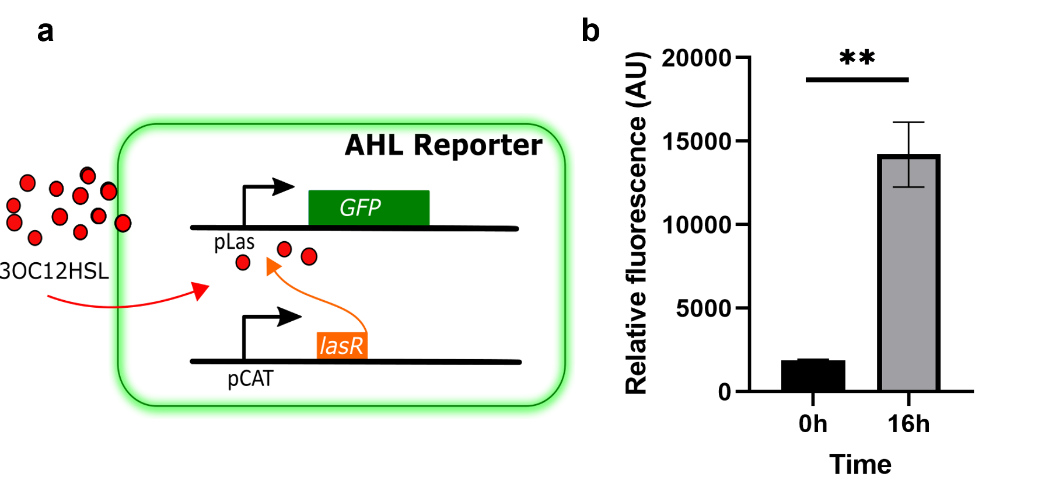

**Figure S7:** *Reporter to detect AHL diffusion.* **a** Schematic of reporter strain, *E. coli* JS006-ALT that expresses *sfGFP* in response to 5 nM AHL molecules. **b** Fluorescence reading from the reporter after 16 h of incubation when induced by 5 nM AHL. n = 3, error bar = s.d. Paired t test, *P* value**.0086

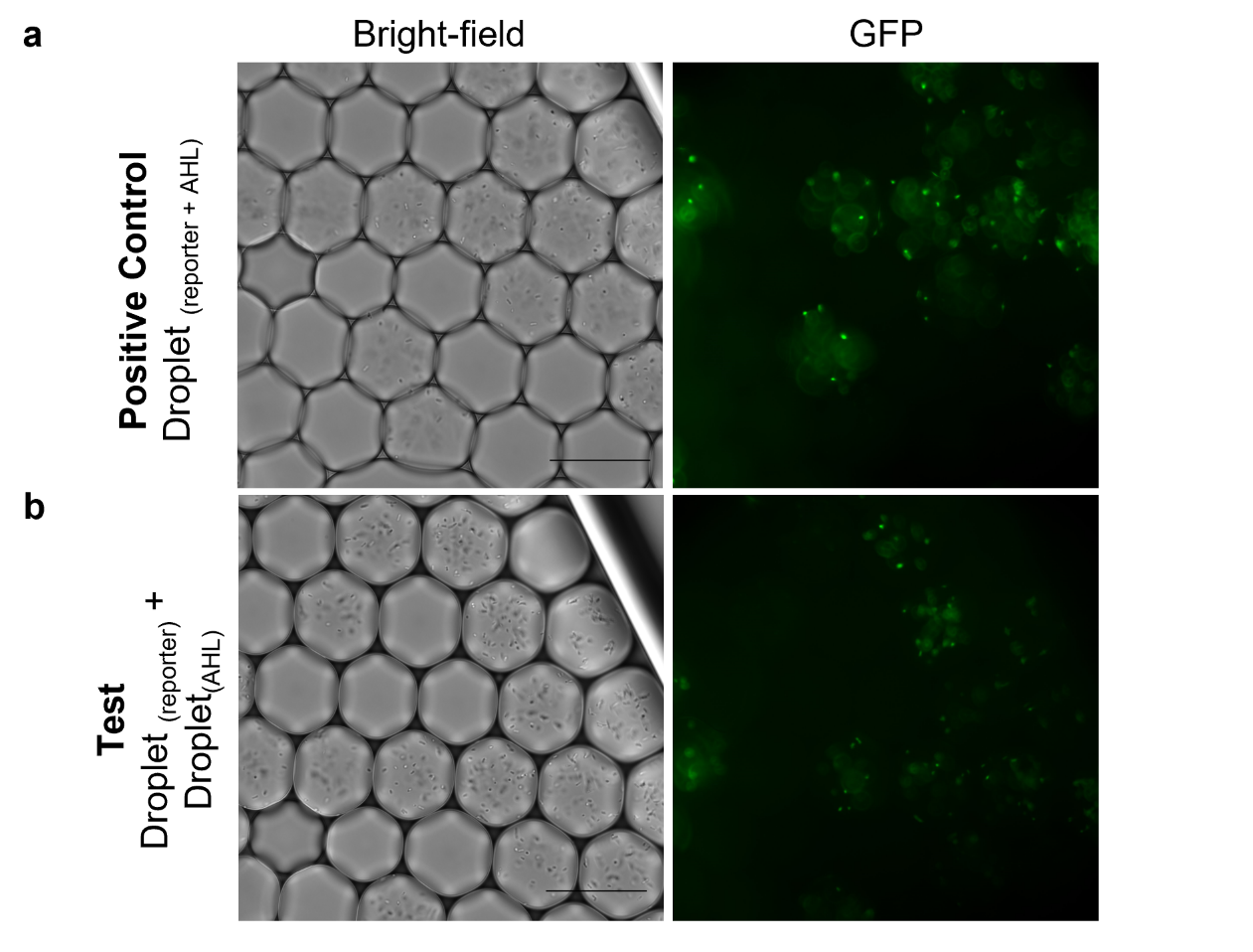

**Figure S8:** *AHL diffuses between droplets in less than 2h.* **a** In the positive controls, 50 nM AHL molecules were added to the same microdroplets as the AHL reporter, whereas in the test condition **(b)**, 50 nM AHL molecules and the reporter cells are encapsulated into different microdroplets that are mixed together. We observe GFP expression by the reporter within 2 hours of incubation. Scale bar 50 µm.

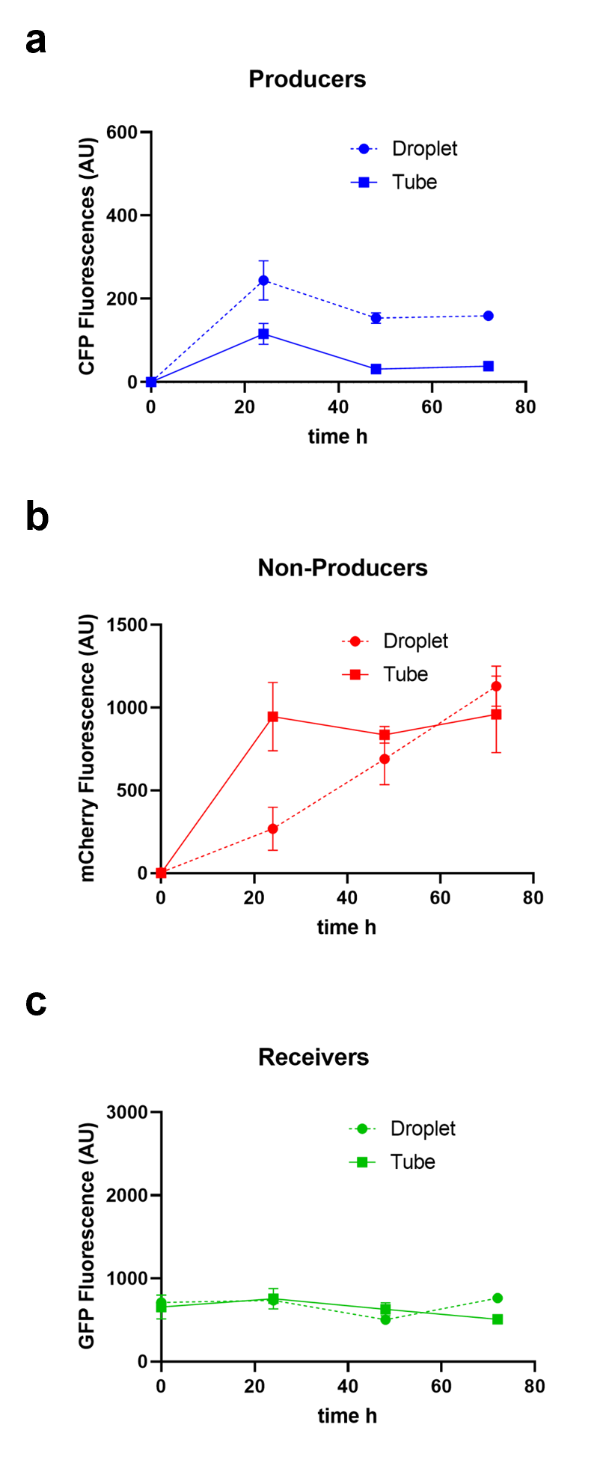

**Figure S9:** *The Producer, the Non-Producer, and the Receiver growth trends in the competition-based experimental strategy.* The fluorescent readings corresponding to Producer **(a)**, Non-Producer **(b)**, and Receiver **(c)** population percentages were used to compare the growth trends of the two culturing conditions (microdroplet = dotted line, test tube = solid line) induced at 1 mM IPTG. The initial population density was chosen as before with $\lambda_{1}= 0.05$ for Producers, $\lambda_{2}=0.05$ for Non-Producer, and $\lambda_{3}=5$ for Receivers. After 24 h of growth, the fluorescence for each population were measured for 3 cycles using Tecan Spark fluorescent microplate reader. n = 2, error bars= s.d.

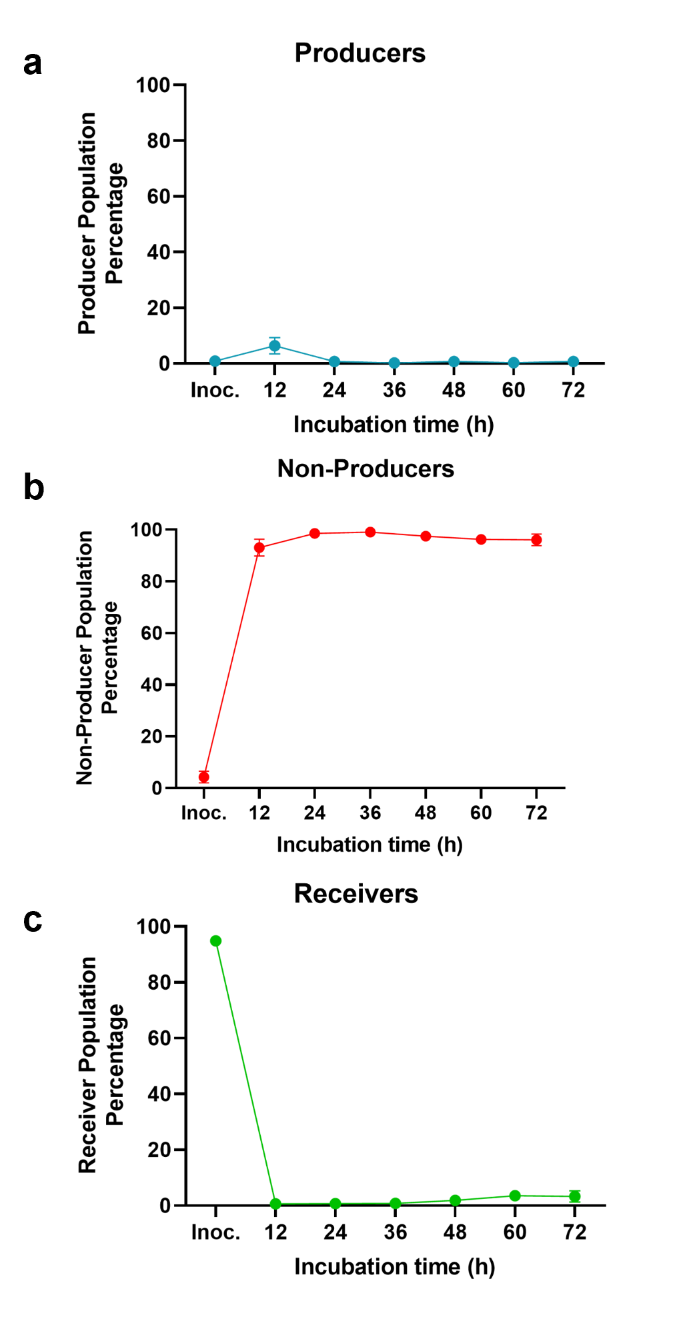

**Figure S10:** *The Producer, the Non-Producer, and the Receiver population percentage for 12 h growth cycles in the competition-based experimental strategy.* The fluorescent readings corresponding to Producer **(a)**, Non-Producer **(b)**, and Receiver **(c)** population percentages were used to report the growth trends for 12 h of growth cycles for each strain. The initial population density was chosen as before with $\lambda_{1}= 0.05$ for Producers, $\lambda_{2}=0.05$ for Non-Producer, and $\lambda_{3}=5$ for Receivers, induced at 1 mM IPTG. After 12 h of growth, the cell counts for each population were measured for 6 cycles using Tecan Spark fluorescent microplate reader. n = 2, error bars= s.d.

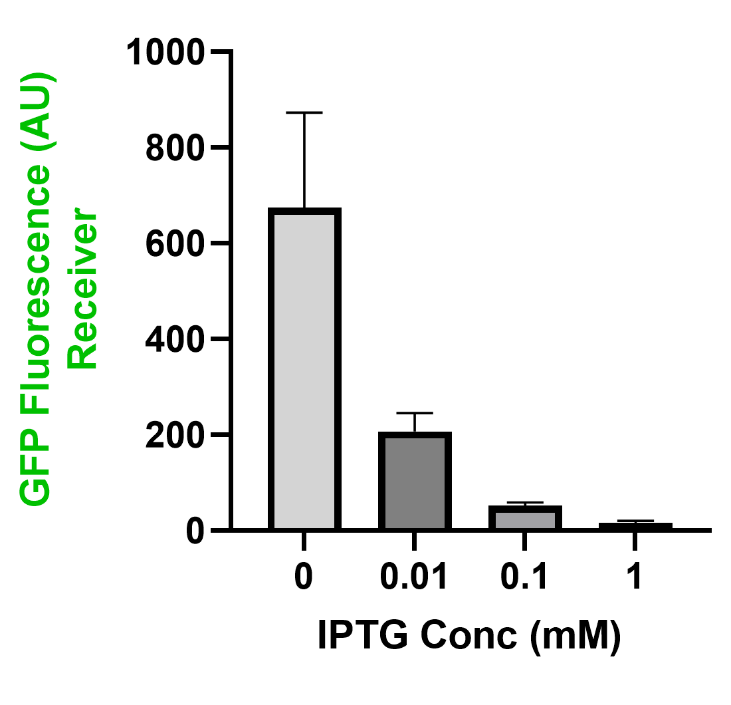
**Figure S****11:** *Receiver killing can be regulated by varying the concentrations of IPTG.* Here, Producers and Receivers are co-cultured in 1:10 population ratio at different IPTG concentrations. At different concentrations of IPTG, Producers secrete different concentrations of AHLs into the culture, which can go on to induce Receiver cell death. As the synthesis of AHL by Producers increases, more Receivers are killed. n = 3, error bars = s.d.

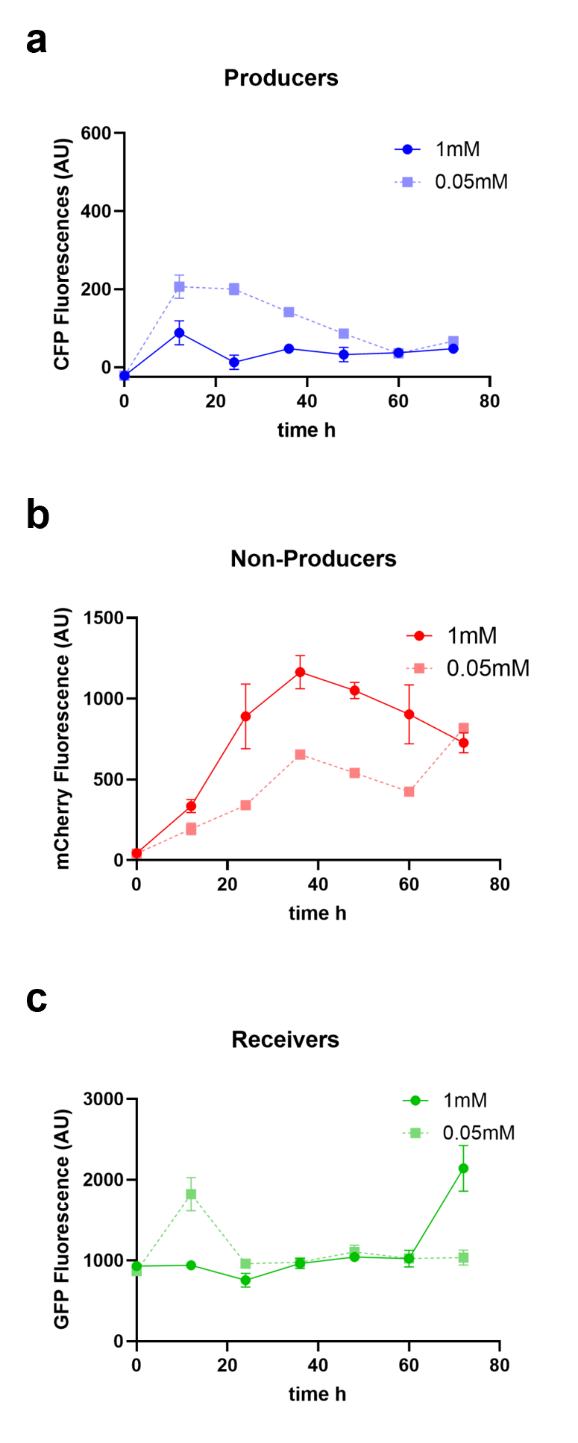

**Figure S12:** *The Producer, the Non-Producer, and the Receiver growth trends induced at different IPTG concentrations in the competition-based experimental strategy.* The fluorescent readings corresponding to Producer **(a)**, Non-Producer **(b)**, and Receiver **(c)** population percentages were used to compare the growth trends of the two culturing conditions (0.05 mM IPTG = dotted line, 1 mM IPTG = solid line). The initial population density was chosen as before with $\lambda_{1}= 0.05$ for Producers, $\lambda_{2}=0.05$ for Non-Producer, and $\lambda_{3}=5$ for Receivers. After 24h of growth, the fluorescence for each population were measured for 3 cycles using Tecan Spark fluorescent microplate reader. n = 2, error bars= s.d.

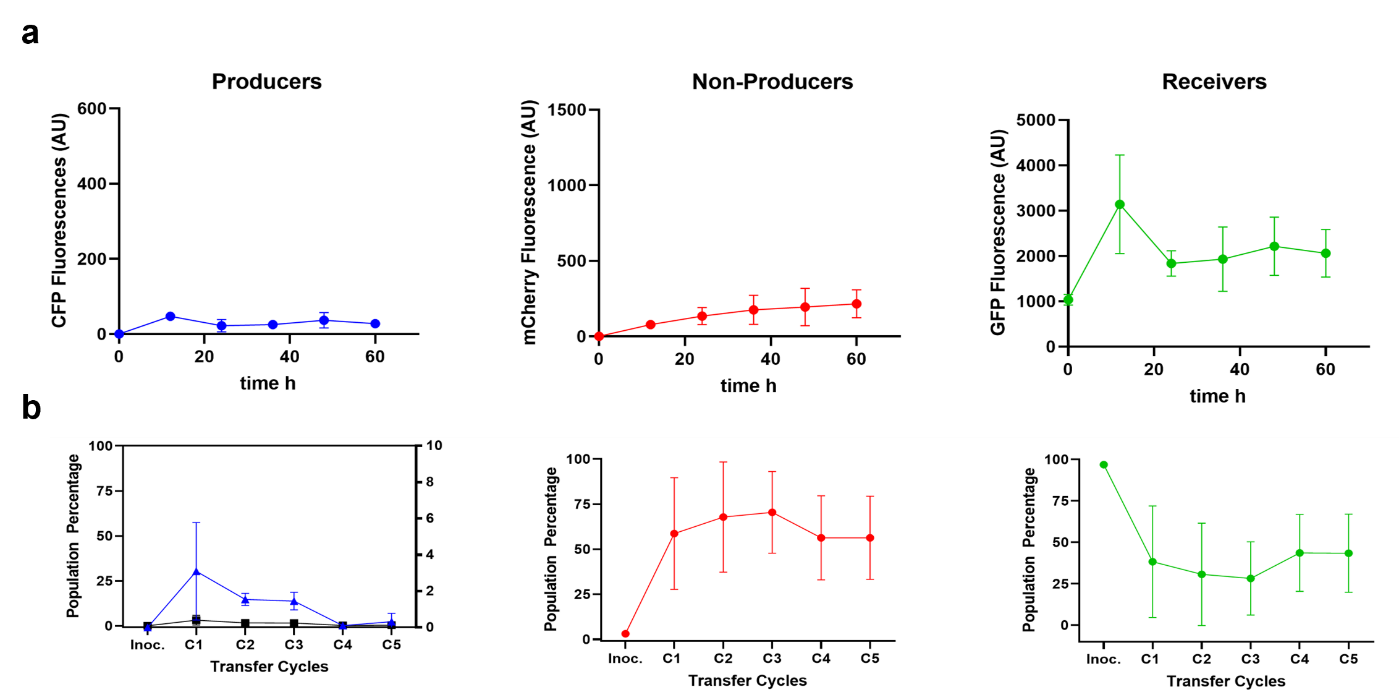
**Figure S13:** *The Producer, Non-Producer, and Receiver growth trends at lower initial Producer population density.* The fluorescence **(a)** and cell counts **(b)** correspond to Producer (blue) , Non-Producer (red), and Receiver (green) population growth trends when induced at 0.05 mM IPTG. The initial population density was chosen as $\lambda_{1}= 0.005$ for Producers (10-folds lower compared to the previous experiments), $\lambda_{2}=0.05$ for Non-Producer, and $\lambda_{3}=5$ for Receivers. After 12 h of growth, the cell counts for each population were measured for 5 cycles. n = 2, error bars = s.d.. The bigger error bars are assumed to be because of the larger variation in the starting Producer population. At lower Producer populations used here, the variability in the initial cell counts are higher when determined using OD values instead of automatic cell counters resulting in higher variability in the interaction of Producers with the other two cell types.

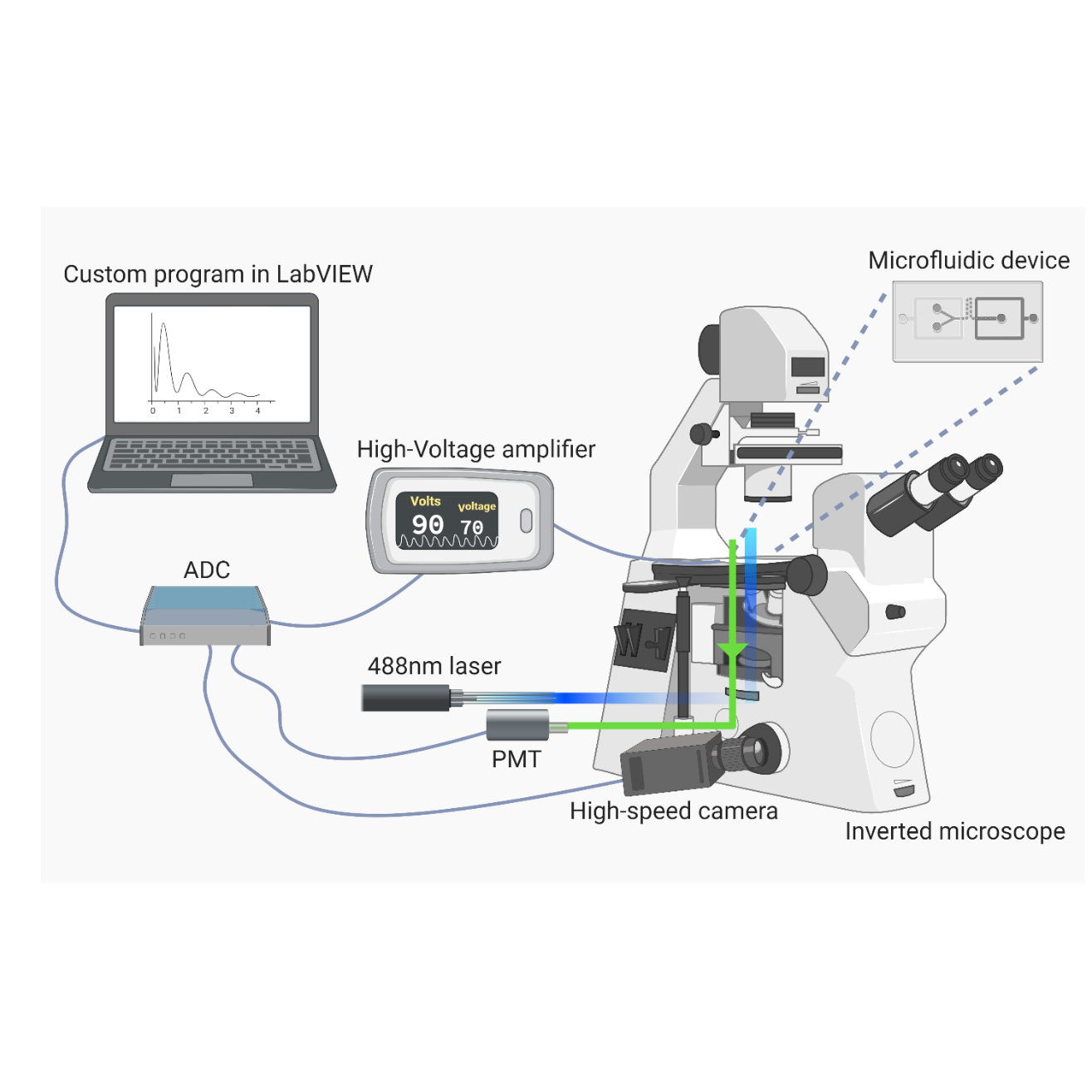

**Figure S14:** *Optical setup for Fluorescent Activated Droplet Sorting (FADS)*. The optical setup includes a laser light (488 nm, 50 mW) directed to a point in the path of droplet flow in microfluidic chip, through an objective of an inverted microscope. The emitted signal from the droplets is processed by two photomultiplier tubes (PMTs), which then transmits the signal to be read by a custom LabVIEW program (Mazutis et al., 2013). When a fluorescent cell is detected, the program activates the electrode through a high-voltage amplifier for sorting the cell into the collection chamber on the device. A custom LabVIEW software developed by Mazutis et al. will be used to record droplet fluorescence and trigger the electrodes during sorting. Data acquisition card (PCIe-7842R; National Instruments) recorded the data at the rate of 100 kHz.

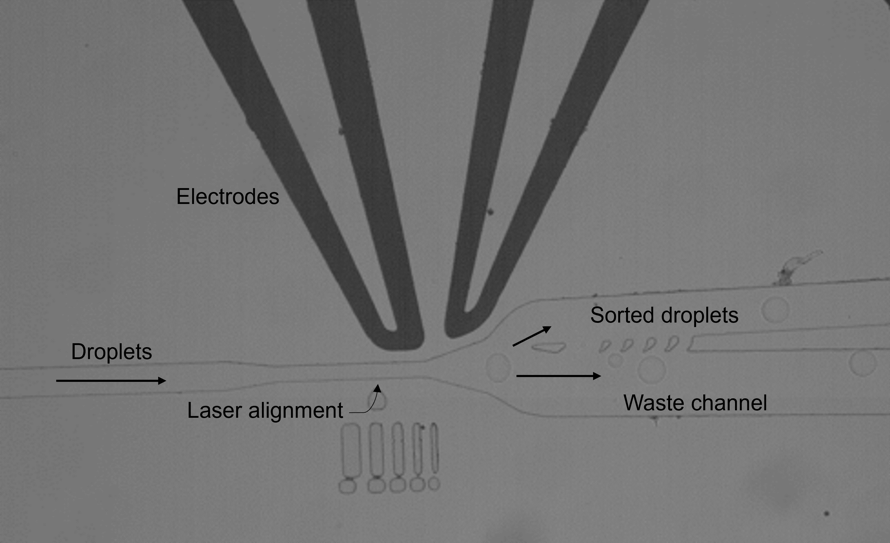

**Figure S15:** *Sorting module of the microfluidic droplet sorter device*. The optimally spaced droplets from the pico-injection module pass through a narrow channel that pinches the droplet for detection. Based on the sorting threshold set on the intensity of the recorded signal, the droplet will be either sorted to the high resistance upper channel by a phenomenon called dielectropheresis or enter the waste channel.

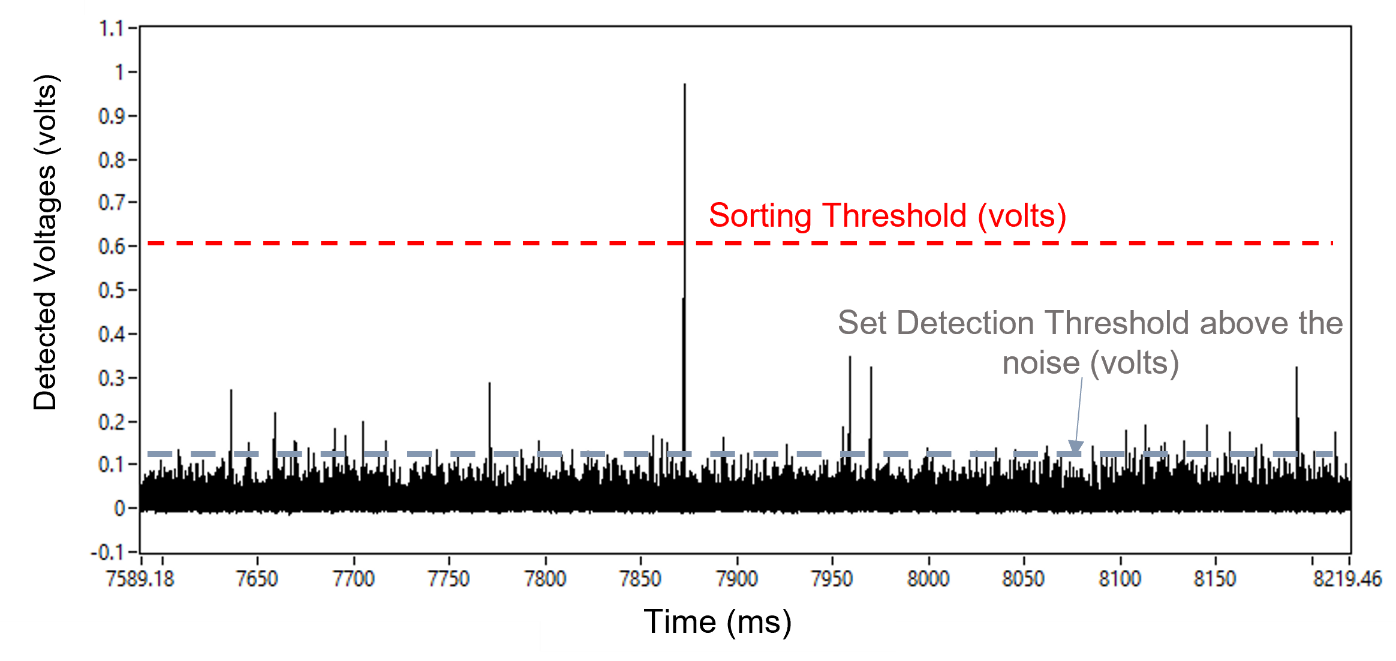

**Figure S16:** *Fluorescence intensity of individual droplets during sorting.* The height of the peak represents the intensity of the detected signal over time. By determining the sorting threshold (voltage amplitude), we can eliminate droplets with no cells or droplets with cells other than GFP-labeled Receiver cells from the collection channel. The peak between 7850 and 7900 ms corresponds to a bright green microdroplet that is targeted for sorting.

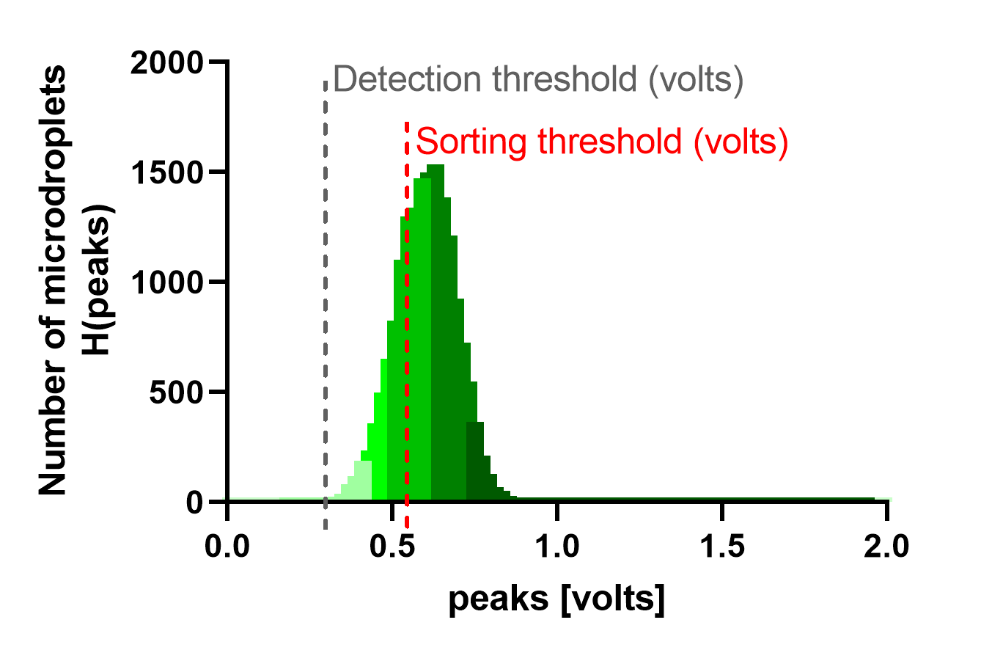

**Figure S17:** *Histogram of the detected microdroplet with green fluorescence intensity.* From the distribution of fluorescent signals of the microdroplet passing through the sorting channel, sorting threshold can be set to sort out microdroplets with green fluorescence signal. Here, the grey dotted line indicates the droplet detection threshold and the red dotted line indicates a sorting threshold in volts. Microdroplets above the sorting threshold will be sorted out. Rest of the microdroplets will be collected off-chip into a microcentrifuge tube for further analysis.

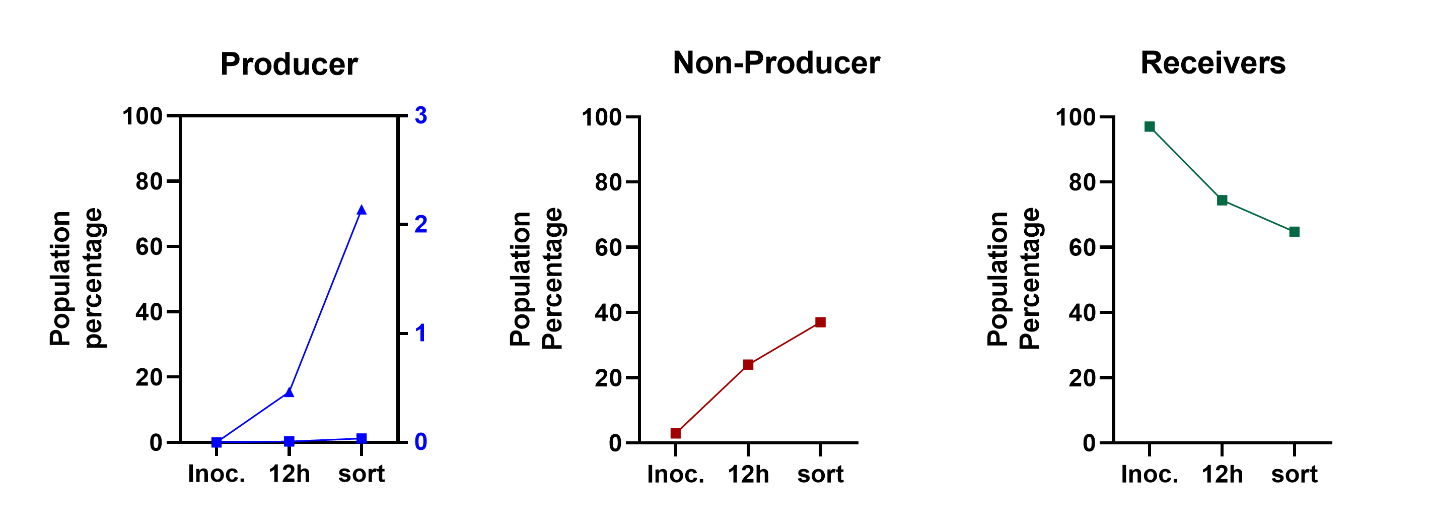

**Figure S18:** Growth trends for the Producers **(a)**, Non-Producers **(b)**, and Receivers **(c)** at inoculation (Inoc.), 12 h of incubation (12h) followed by the fluorescent sorting (sort) at 85% thresholds. Total number of droplets screened, n = ~120k microdroplets.

**Model Supplement**

**1 Population model in well-mixed co-cultures**

Let $P\left( t \right), N\left( t \right),$ and $R(t)$ represent the population count of the Producers, Non-Producers, and Receivers, with growth rate $g_{i}$, $i=1,2,3$ respectively, at time $t$. Assume cells are growing in a well-mixed environment, where diffusive signals are uniformly distributed. Let $Q(t)$ denote the number of AHL signaling molecules, which are being synthesized by Producers at some constant rate $g_{4}$ and have degradation rate $\gamma$. Let $T$ represent the incubation cycle length. Assuming logistic growth with a carrying capacity of $S_{max}$, we then have the following ODE model for a single incubation cycle in $t\in[0,T]$,

$$\begin{aligned} \dot{P}=g_{1} \left( 1-{S(t)}/{S_{\max}} \right)P(t), \#\left( S1a \right) \end{aligned}$$

$$\begin{aligned} \dot{N}=g_{2}\left( 1-{S(t)}/{S_{\max}} \right)N(t),\#\left( S1b \right) \end{aligned}$$

$$\begin{aligned} \dot{R}=g_{3} \left( 1-{S\left( t \right)}/{S_{\max}} \right)R(t)-\alpha f\left( Q(t) \right)R(t),\#\left( S1c \right) \end{aligned}$$

$$\begin{aligned} \dot{Q}=g_{4}P(t)-\gamma Q(t).\#\left( S1d \right) \end{aligned}$$

Here, the AHL-induced lysis rate is modeled using a standard Hill function $f$ of the AHL signal concentration $Q\left( t \right)$ with threshold $\theta$ and Hill coefficient $m$, where

$$\begin{aligned} f\left( Q\left( t \right) \right)=\frac{{Q\left( t \right)}^{m}}{\theta^{m}+{Q\left( t \right)}^{m}}, \end{aligned}$$

with maximum lysis rate of $\alpha$. Moreover, we represent the total resource consumed using a weighted sum $S\left( t \right)=wP\left( t \right)+N\left( t \right)+R\left( t \right)$. The weight $w\geq1$ accounts for the fitness cost of Producer cells synthesizing AHL molecules, where $w$ increases with [IPTG]. In particular, the value of $k$ is chosen to match the experimentally observed population densities, as shown in **Fig. 1d.** For low inducer levels, such as [IPTG] $\leq0.1$ mM, we set $w=1$. For [IPTG]$=1$mM, we set $w=1.3/0.8\approx1.6$.

Since the cell counts for the well-mixed co-culture are in the millions as shown in **Table S2**, we normalize all variables by ${10}^{6}$ when simulating equations (S1). We also consider a normalized production rate $g_{4}$ with respect to the threshold by setting $\theta=1\times{10}^{6}$. Moreover, since AHL signal degradation is negligible over the course of each incubation cycle^1^, we set $\gamma=0$.

When iterating the incubation cycles, the co-culture is diluted $d$-fold before entering the next round of incubation. After dilution, washed cells are transferred into fresh media, therefore we assume the initial AHL signal level in the extracellular environment is 0 for the next cycle. The iterative cycle can be modeled using the following chart.

$$\left[ P\left( 0 \right), N\left( 0 \right),R\left( 0 \right), Q\left( 0 \right) \right]^{T}\underset{\to}{Eq. \left( S1 \right)}\left[ P\left( T \right), N\left( T \right),R\left( T \right), Q\left( T \right) \right]^{T}\underset{\to}{\mathrm{Dilution}}\left[ P\left( T \right), N\left( T \right),R\left( T \right), 0 \right]^{T}/d$$

$$\mathrm{iteration}$$

**2 Population model in the microdroplet system**

When incubating the cells in the microdroplet system, we need to extend the above model to incorporate the stochastic nature of the co-encapsulation and the spatial segregation of the growth environment. While it’s computationally expensive to simulate each of the millions of microdroplets, we capture the essence of the microdroplet system by studying the distribution of various combination of cells initially loaded in the microdroplets. We then characterize the microdroplets by their initial conditions and calculate the total population as a weighted sum of those different microdroplets.

**2.1 Stochastic Co-encapsulation**

Given a well-mixed single-strain culture, let $\lambda$ denote the expected number of cells per microdroplet. We assume the probability of a microdroplet containing $n$ cells follows a Poisson distribution{Citation}^2^, where the probability mass function $g$ is defined as

$$\begin{aligned} g\left( n;\lambda\right)=\frac{e^{-\lambda}\lambda^{n}}{n!}. \end{aligned}$$

From there, we get that the probability of a given microdroplet being non-empty is

$$\begin{aligned} \Pr\left( n>0 \right)=1-g\left( 0;\lambda\right)=1-e^{-\lambda}. \end{aligned}$$

Next, we assume the number of cells from each strain encapsulated into a microdroplet is independent from that of other strains. Let $\lambda_{i}$ represents the average number of cells per microdroplet for the corresponding strain. We then have that the probability of a microdroplet containing $n_{i}$ cells of strain $i$, for $i=1,2,\ldots$, can calculated as^3^

$$\begin{aligned} \Pr\left( n_{1},n_{2},\ldots\right) =\prod_{i} g\left( n_{i};\lambda_{i} \right)=\prod_{i} \frac{e^{-\lambda_{i}}\lambda_{i}^{n_{i}}}{n_{i}!}.\#\left( S2 \right) \end{aligned}$$

In our case, when co-encapsulating Producers ($\lambda_{1}=0.1$) with Receivers ($\lambda_{2}=5$), we find that the probability of a microdroplet contains only Receivers (R-only microdroplets) is

$$\begin{aligned} \Pr\left( n_{1}=0,n_{2}>0 \right) =e^{-\lambda_{1}}\left( 1-e^{-\lambda_{2}} \right)\approx89.87\%. \#\left( S3 \right) \end{aligned}$$

After co-encapsulation, each microdroplet can be characterized by a vector $\boldsymbol{n}=\left( n_{1},n_{2},n_{3} \right)^{T}$, whose components describe the enclosed cell count for each strain. For each initial condition $\boldsymbol{n}$, we can calculate the likelihood of such a microdroplet being made by using equation $\left( S2 \right)$.

**2.2 Intra- and Inter-microdroplet interactions**

Let $M$ denotes the total number of microdroplets made. In each microdroplet, assume cell growth follows the logistic model with carrying capacity $\hat{S}_{\max}$. Let $P_{i}\left( t \right), N_{i}\left( t \right),$ and $R_{i}(t)$ represent the populations of Producers, Non-Producers, and Receivers, respectively, in the $i^{th}$ microdroplet at time $t$. Assume that there are $n_{1i}, n_{2i}, n_{3i}$ number of cells initially enclosed in microdroplet $i$ from each strain. For each incubation cycle $t\in[0,T]$, we have that population dynamics in the $i^{th}$ microdroplet are governed by the following equations.

$$\begin{aligned} \dot{P}_{i}=g_{1} \left( 1-{S_{i}(t)}/{\hat{S}_{\max}} \right)P_{i}(t), \#\left( S4a \right) \end{aligned}$$

$$\begin{aligned} \dot{N}_{i}=g_{2}\left( 1-{S_{i}(t)}/{\hat{S}_{\max}} \right)N_{i}(t),\#\left( S4b \right) \end{aligned}$$

$$\begin{aligned} \dot{R}_{i}=g_{3} \left( 1-{S_{i}(t)}/{\hat{S}_{\max}} \right)R-f\left( Q_{i}(t) \right)R_{i}(t),\#\left( S4c \right) \end{aligned}$$

where $S_{i}\left( t \right)=wP_{i}\left( t \right)+N_{i}\left( t \right)+R_{i}\left( t \right)$ and $Q_{i}(t)$ represents the effective AHL signal level in the $i^{th}$ microdroplet.

The dynamics of each microdroplet are coupled if AHL signals diffuse among microdroplets. Let parameter $\beta$ denote the percentage of the signal produced that stays within the microdroplet, with the other $1-\beta$ percentage of the signal being immediately shared with all microdroplets. We then have the signal level in the $i^{th}$ microdroplet, $Q_{i}(t)$, satisfies the following equation,

$$\begin{aligned} \dot{Q}_{i}=\beta g_{4}P_{i}(t)+\left( 1-\beta\right)\frac{g_{4}}{M}\sum_{k=1}^{M} P_{k}(t)-\gamma Q_{i}(t).\#\left( S5 \right) \end{aligned}$$

**2.3 Model Reduction**

Assume that microdroplet $i$ and microdroplet $\hat{i}$ have the same initial conditions. That is, the two microdroplets contain the same number of Producer, Non-Producer, and Receiver cells after the initial co-encapsulation. By the uniqueness of the solution for equations (S4-S5), we have that microdroplet $i$ and $\hat{i}$ have the same solution, where

$$X_{i}\left( t \right)=X_{\hat{i}}\left( t \right),$$

for $X=P,N,R,Q$. This implies that the total population of each strain in the system can be represented as a weighted sum of the sub-populations in microdroplets with different types of initial conditions.

The above reduction allows us to only track microdroplets with distinct initial conditions. If the initial conditions for all microdroplets are known, the weights of each type of initial condition are equal to their frequency. In this case, due to technical limitations, we can’t count the number of cells from each strain inside a microdroplet. Instead, we estimate the frequency of different initial conditions using equation (S2).

Let $j$ index different types of initial conditions with $\boldsymbol{n}_{j}=\left( n_{1j}, n_{2j}, n_{3j} \right)^{T}$. Let $p_{j}$ represent the probability of a microdroplet being made containing $n_{1j}$ Producers, $n_{2j}$ Non-Producers, and $n_{3j}$ Receivers. We refer to the microdroplets with an initial condition of $\boldsymbol{n}_{j}$ as the $j^{th}$ type of microdroplet. For computational efficiency, we only consider initial conditions with expected frequency $p_{j}=\Pi_{i=1}^{3}g\left( n_{ij};\lambda_{i} \right)$ higher than a chosen threshold $\epsilon$. For example, when $\epsilon$ is set to be ${10}^{-6}$, that means we don’t consider microdroplets with an initial condition less likely than 1 in 1 million. See **Table S3** for an example of the 20 most likely microdroplets types for the given $\lambda_{i}$ values.

We now have the following model. Let $\hat{M}$ denotes the number of distinct initial conditions that have probability greater than $\epsilon$. Let the matrix $A=\left[ \boldsymbol{n}_{j} \right]_{3\times\hat{M}}$ represent the initial condition space. The column vectors $\boldsymbol{n}_{j}=\left( n_{1j}, n_{2j}, n_{3j} \right)^{T}$ satisfy

$$\boldsymbol{n}_{j}\in\left\{ \left( n_{1}, n_{2}, n_{3} \right)^{T}\in\mathbb{Z}_{\geq0}^{3\times1} \right|\Pi_{i=1}^{3}g\left( n_{i};\lambda_{i} \right)\geq\epsilon\}.$$

We rank the initial conditions by their probabilities. That is, $p_{j}\geq p_{\hat{j}}$ if $j\geq\hat{j}$.

For the $j^{th}$ type of microdroplet with initial condition $\left( P_{j}\left( 0 \right),N_{j}\left( 0 \right),R_{j}\left( 0 \right) \right)^{T}=\boldsymbol{n}_{j}$ and $Q_{j}\left( 0 \right)=0$, the population dynamics can be described by the following system,

$$\begin{aligned} \dot{P}_{j}=g_{1} \left( 1-{S_{j}(t)}/{\hat{S}_{\max}} \right)P_{j}(t), \#\left( S6a \right) \end{aligned}$$

$$\begin{aligned} \dot{N}_{j}=g_{2}\left( 1-{S_{j}(t)}/{\hat{S}_{\max}} \right)N_{j}(t),\#\left( S6b \right) \end{aligned}$$

$$\begin{aligned} \dot{R}_{j}=g_{3} \left( 1-{S_{j}(t)}/{\hat{S}_{\max}} \right)R(t)-f\left( Q_{j}(t) \right)R_{j}(t),\#\left( S6c \right) \end{aligned}$$

$\begin{aligned} \dot{Q}_{j}=\beta g_{4}P_{j}(t)+\left( 1-\beta\right)g_{4}\sum_{k=1}^{\hat{M}} {p_{k}P}_{k}(t)-\gamma Q_{j}(t).\#\left( S6d \right) \end{aligned}$

with $S_{j}\left( t \right)=wP_{j}\left( t \right)+N_{j}\left( t \right)+R_{j}\left( t \right)$. The total population for each strain $X=P,N,R$ can be calculated as the weighted sum

$$\begin{aligned} X\left( t \right)=M\sum_{k=1}^{\hat{M}} {p_{k}X}_{k}\left( t \right). \end{aligned}$$

At the end of an incubation cycle, $X\left( T \right)$ gives us the final population for each strain. After the $d$-fold dilution of the resulting culture and replacing of the media, new microdroplets are made with $\boldsymbol{\lambda}=\left( \lambda_{1},\lambda_{2},\lambda_{3} \right)^{T}$ set to $X\left( T \right)/d$, which then changes the probability vector $\boldsymbol{p=}\left( p_{1}, \ldots,p_{j},\ldots, p_{\hat{M}} \right)^{T}$ for the next cycle. The iterative model is presented by the following chart.

$$\left[ \begin{aligned} P\left( 0 \right) \\ N\left( 0 \right) \\ R\left( 0 \right) \\ Q\left( 0 \right) \end{aligned} \right]\underset{\begin{aligned} \boldsymbol{\lambda}={X\left( 0 \right)}/M \\ p_{j}=\Pi_{i}g\left( n_{ij};\lambda_{i} \right) \end{aligned}}{\underset{\to}{Co-encapsulation}}\left[ \ldots\begin{aligned} P_{j}\left( 0 \right) \\ N_{j}\left( 0 \right) \\ R_{j}\left( 0 \right) \\ Q_{j}\left( 0 \right) \end{aligned} \ldots\right]\left[ \begin{aligned} \begin{aligned} p_{1} \\ \vdots\\ p_{j} \\ \vdots\end{aligned} \\ p_{\hat{M}} \end{aligned} \right]\underset{\to}{Eq. \left( S6 \right)}\left[ \ldots\begin{aligned} P_{j}\left( T \right) \\ N_{j}\left( T \right) \\ R_{j}\left( T \right) \\ Q_{j}\left( T \right) \end{aligned} \ldots\right]\left[ \begin{aligned} \begin{aligned} p_{1} \\ \vdots\\ p_{j} \\ \vdots\end{aligned} \\ p_{\hat{M}} \end{aligned} \right]\underset{\mathrm{Refill}}{\underset{\to}{\mathrm{Dilution}}} \left[ \begin{aligned} {P\left( T \right)}/d \\ {N\left( T \right)}/d \\ {R\left( T \right)}/d+R_{0} \\ 0 \end{aligned} \right]$$

$$\mathrm{iteration}$$

**2.4 Sorting**

In our experimental design, sorting the microdroplets is based on the fluorescent intensity from the GFP-tagged Receiver cells within each microdroplet. Assume that GFP is produced by the Receiver cells at a constant rate $g_{5}$ and being degraded at rate $\gamma_{f}$. Let $F_{j}(t)$ represents the GFP levels in the $j^{th}$ type of microdroplet. Then we have that

$$\begin{aligned} \dot{F_{j}}=g_{5}R_{j}\left( t \right)-\gamma_{f}F_{j}\left( t \right). \#\left( S13 \right) \end{aligned}$$

Since GFP degrade slowly^4^ over the course of each incubation cycle, we set $\gamma_{f}=0$. Moreover, since GFP level is a relative measurement, we consider a normalized model by setting $g_{5}=1$.

Assume the sorting threshold is set to be $\mu\in[0\%,100\%]$. At the end of the incubation cycle before dilution, we will rank the microdroplets using $F_{j}\left( T \right)$ and sort out the top $\mu$ portion of all microdroplets. In the simulation, this is implemented by updating the weight of each type of droplet as demonstrated in **Table S3**.

**Table S1:** Growth rate of the three strains at different IPTG concentrations **(Fig. 1d)**

| Strain Type/ IPTG Concentrations | 0 mM IPTG | 0.1 mM IPTG | 1 mM IPTG |
| --- | --- | --- | --- |
| Producers (P) | 0.5873 /h | 0.6155 /h | 0.5404 /h |
| Non-Producers (N) | 0.6327 /h | 0.6613 /h | 0.6416 /h |
| Receivers (R) | 0.5235 /h | 0.5067 /h | 0.5090 /h |

**Table S2:** Parameter values used for simulations in **Fig. 3e, 4b, 5c-d, and S3**.

| **Name** | **Description** | | **Value** | **Unit** | | **Reference** |
| --- | --- | --- | --- | --- | --- | --- |
| Initial Co-culture for Well-mixed Test Tube Condition | | | | | | |
| $\rho_{1}$ | Producer Density | | $1.5$ | $\times{10}^{6}/\mathrm{ml}$ | | Experimental Data |
| $\rho_{2}$ | Non-Producer Density | | $1.5$ | $\times{10}^{6}/\mathrm{ml}$ | |  |
| $\rho_{3}$ | Receiver Density | | $1.5$ | $\times{10}^{8}/\mathrm{ml}$ | |  |
| $V$ | Co-culture Volume | | $5$ | $\mathrm{ml}$ | |  |
| Initial Co-culture and Co-encapsulation for Microdroplet Condition | | | | | | |
| $\rho_{1}$ | Producer Density | | $1.5/0.015$ | $\times{10}^{6}/\mathrm{ml}$ | | Experimental Data |
| $\rho_{2}$ | Non-Producer Density | | $1.5$ | $\times{10}^{6}/\mathrm{ml}$ | |  |
| $\rho_{3}$ | Receiver Density | | $1.5$ | $\times{10}^{8}/\mathrm{ml}$ | |  |
| $V$ | Co-culture Volume | | $0.5$ | $\mathrm{ml}$ | |  |
| $v$ | Microdroplet Volume | | $33.5$ | $\mathrm{pl}$ | |  |
| Population Growth | | | | | | |
| $g_{1}$ | Producer Growth Rate | 1 mM IPTG | $0.5404$ | $h^{-1}$ | | **Table S1** |
|  |  | 0.05 mM IPTG | $0.6078$ | $h^{-1}$ | |  |
| $g_{2}$ | Non-Producer Growth Rate | | $0.6452$ | $h^{-1}$ | | Estimated from **Table S1** |
| $g_{3}$ | Receiver Growth Rate | | $0.5131$ | $h^{-1}$ | |  |
| $S_{\max}$ | Carrying Capacity | Tube | $2$ | $\times{10}^{9}$ | | Experimental Data |
|  |  | Microdroplet | $150$ |  | |  |
| $w$ | Producer Growth Weight | 1 mM IPTG | $1.6$ | | | Estimated with data from **Fig. 1d** |
|  |  | 0.05 mM IPTG | $1$ | | |  |
| AHL-induced Lysis of Receivers | | | | | | |
| $g_{4}$ | Production Rate of AHL | 1 mM IPTG | $10$ | $h^{-1}$ | | Estimated from Experimental Data |
|  |  | 0.05 mM IPTG | $0.5$ | $h^{-1}$ | |  |
| $\alpha$ | Maximum Lysis Rate | | $0.65$ | $h^{-1}$ | |  |
| $\theta$ | AHL Threshold of Lysis | | ${V\times10}^{6}ml^{-1}$ | | |  |
| $m$ | Hill Coefficient | | $2$ | | |  |
| Iterative Incubation Cycle | | | | | | |
| $T$ | Incubation Time per Cycle | | $24/12$ | | $h$ | Experimental Data |
| $d$ | Dilution Factor | | $50$ | | |  |

**Table S3:** The first 20 most likely droplet types, ranked by their expected percentages from co-encapsulation, and changes to their percentages after sorting at different thresholds, corresponding to simulation results from **Fig. 5**. The initial percentages of each droplet type are estimated from equation (S2), with $\lambda_{1}=5\times{10}^{-4}$ (P)*,* $\lambda_{2}=5\times{10}^{-2}$ (N), $\lambda_{3}=5$(R). After sorting, the $p_{j}=0$ indicates that all droplets of type $j$ are sorted out.

| Droplet Type | Cells Encapsulated | | | Percentages of Droplet Type $j$, $p_{j}$ | | | |
| --- | --- | --- | --- | --- | --- | --- | --- |
|  | P | N | R | Initially Generated | After Sorting at Threshold | | |
| $j$ | $n_{1j}$ | $n_{2j}$ | $n_{3j}$ |  | 75% | 85% | 95% |
| 1 | 0 | 0 | 5 | 16.68% | 0 | 0 | 0 |
| 2 | 0 | 0 | 4 | 16.59% | 0 | 0 | 0 |
| 3 | 0 | 0 | 6 | 13.97% | 0 | 0 | 0 |
| 4 | 0 | 0 | 3 | 13.21% | 10.29% | 0.29% | 0 |
| 5 | 0 | 0 | 7 | 10.03% | 0 | 0 | 0 |
| 6 | 0 | 0 | 2 | 7.89% | 7.89% | 7.89% | 0 |
| 7 | 0 | 0 | 8 | 6.30% | 0 | 0 | 0 |
| 8 | 0 | 0 | 9 | 3.52% | 0 | 0 | 0 |
| 9 | 0 | 0 | 1 | 3.14% | 3.14% | 3.14% | 3.14% |
| 10 | 0 | 0 | 10 | 1.77% | 0 | 0 | 0 |
| 11 | 0 | 1 | 5 | 0.84% | 0.84% | 0.84% | 0 |
| 12 | 0 | 1 | 4 | 0.83% | 0.83% | 0.83% | 0 |
| 13 | 0 | 0 | 11 | 0.81% | 0 | 0 | 0 |
| 14 | 0 | 1 | 6 | 0.70% | 0 | 0 | 0 |
| 15 | 0 | 1 | 3 | 0.66% | 0.66% | 0.66% | 0.57% |
| 16 | 0 | 0 | 0 | 0.62% | 0.62% | 0.62% | 0.62% |
| 17 | 0 | 1 | 7 | 0.50% | 0 | 0 | 0 |
| 18 | 0 | 1 | 2 | 0.40% | 0.40% | 0.40% | 0.40% |
| 19 | 0 | 0 | 12 | 0.34% | 0 | 0 | 0 |
| 20 | 0 | 1 | 8 | 0.32% | 0 | 0 | 0 |
